# A single-nucleus multiome analysis of transcriptome and chromatin accessibility reveals cell-type-specific immune modulation for chronic cannabis use among people with HIV infection

**DOI:** 10.64898/2026.02.26.708082

**Authors:** Mingrui Li, Kesava Asam, Xiaoke Duan, Grier P. Page, Ying Hu, Claudia Martinez, Mardge H. Cohen, Nancie M. Archin, Amir Valizadeh, Dana B. Hancock, Eric Johnson, Bradley E Aouizerat, Ke Xu

**Author notes:** **Corresponding address to: Ke Xu, MD, PhD** Professor of Psychiatry Yale School of Medicine Connecticut VA Healthcare, Biomedical Informatics and Data Science 300 George st, Suite 901, New Haven, CT 06511.

## Abstract

As cannabis use continues to rise among people with HIV (PWH), understanding its impact on immune function in this population is becoming increasingly important. To provide new insights on how cannabis modulates immune function, we analyzed single-nucleus multi-omic profiles of peripheral blood mononuclear cells (PBMCs) from PWH to characterize the changes in gene expression and chromatin accessibility associated with chronic cannabis exposure. We identified numerous differentially expressed genes (DEGs) between cannabis users and non-users in each cell type, approximately half of which were unique to individual cell types. Changes in pro- and anti-inflammatory gene expression associated with cannabis use are dependent on cell lineage and type. We identified hundreds of differential chromatin accessibility regions in each cell type, including *cis*-regulatory elements correlated with cell-type-dependent DEGs (e.g. *NFKBIA* in CD4+ T cells and *CCL3L1* in classical monocytes). Multiple cannabis-associated transcription factors (e.g., *NFKB1, FOS*, and *TCF7*) emerge as regulators of the differentially expressed inflammatory genes. Furthermore, cannabis altered the communication between classical monocytes and lymphocytes. These findings indicate that cannabis-induced immunomodulatory effects are profound, dynamic and complex among cell types and that transcriptional changes are regulated at least in part by epigenetic mechanisms.

## Introduction

Cannabis is one of the most widely used drugs in the world^1^, particularly among people with HIV (PWH) infection. The prevalence of cannabis use is as high as 77% among PWH^2^. Chronic HIV infection is characterized by persistent immune activation and heightened inflammation, which significantly increases the risk of comorbidities and mortality among PWH^3–6^. Some studies suggested that cannabis could alleviate the heightened inflammatory state observed in PWH^7–9^, garnering interest in its use as an adjunct to antiretroviral therapy. However, the evidence regarding the benefits of cannabinoids remains inconsistent or contradictory^10,11^. The mechanisms of the cannabis’ immunomodulatory effects in PWH have not been fully understood.

Cannabis’ immunomodulation has long been of interest because of the critical role of its cognate target, cannabinoid receptor 2 (CNR2), in immune modulation and inflammation^12,13^. *CNR2* is expressed in various immune cells, including T and B lymphocytes, natural killer (NK) cells, monocytes, and neutrophils^12–15^, with the highest expression in B cells. More recently, other proteins (i.e., GPR18, GPR55) were demonstrated to act as cannabinoid receptors in immune cells^16–18^. Activation of these receptors has been shown to induce cellular apoptosis^19,20^, inhibit pro-inflammatory cytokine expression^21–24^, and shift the immune response from Th1 to Th2^25,26^.

Much of the existing research has either focused on *in vitro* studies or involved studies employing HIV-uninfected cells, providing limited insight into the real-world complexity of cannabis’ effects on the immune system in the context of HIV infection. Chronic HIV infection could be a powerful model to study cannabis’ immunomodulatory effect because of the persistent low-degree inflammatory state experienced by PWH even among persons who are virologically suppressed (i.e., HIV viral RNA below the limit of detection in blood). In addition, the gene regulatory effects of cannabis exposure have primarily been studied in bulk tissue^27–30^, with little understanding in its cell-type-specific effects. Single-nucleus (sn) multiomic analysis can elucidate the critical roles of gene expression and chromatin accessibility at the cellular level resulting from chronic cannabis exposure.

We previously reported a single cell transcriptome analysis of an *in vivo* acute THC exposure, a principal component of cannabis, on PBMCs in people without HIV infection^31^. In the present study, we aimed to better understand the impacts of chronic cannabis exposure on the immune system and its underlying epigenetic mechanisms in the context of chronic HIV infection. Single-nucleus RNA sequencing (snRNA-seq) and single-nucleus assay for transposase-accessible chromatin with sequencing (snATAC-seq) were performed on the same nuclei to profile transcriptomes and chromatin accessibility of PBMCs from virally suppressed PWH receiving antiretroviral therapy. This cohort included clinically matched cannabis users (CBUs) and non-users (non-CBUs). Here, we characterized cell-type-specific differentially expressed genes (DEGs) and differentially accessible regions (DARs) between CBUs and non-CBUs as well as the *cis*-regulatory architecture linking DARs to DEGs. Cell-type-specific transcription factor (TF) network analysis identified cannabis-relevant TFs and their target genes. Finally, cell-cell communication inferred from transcriptomic profiles was compared between CBUs and non-CBUs. The analytical strategy is presented in **Figure 1A**.

**Figure 1.**
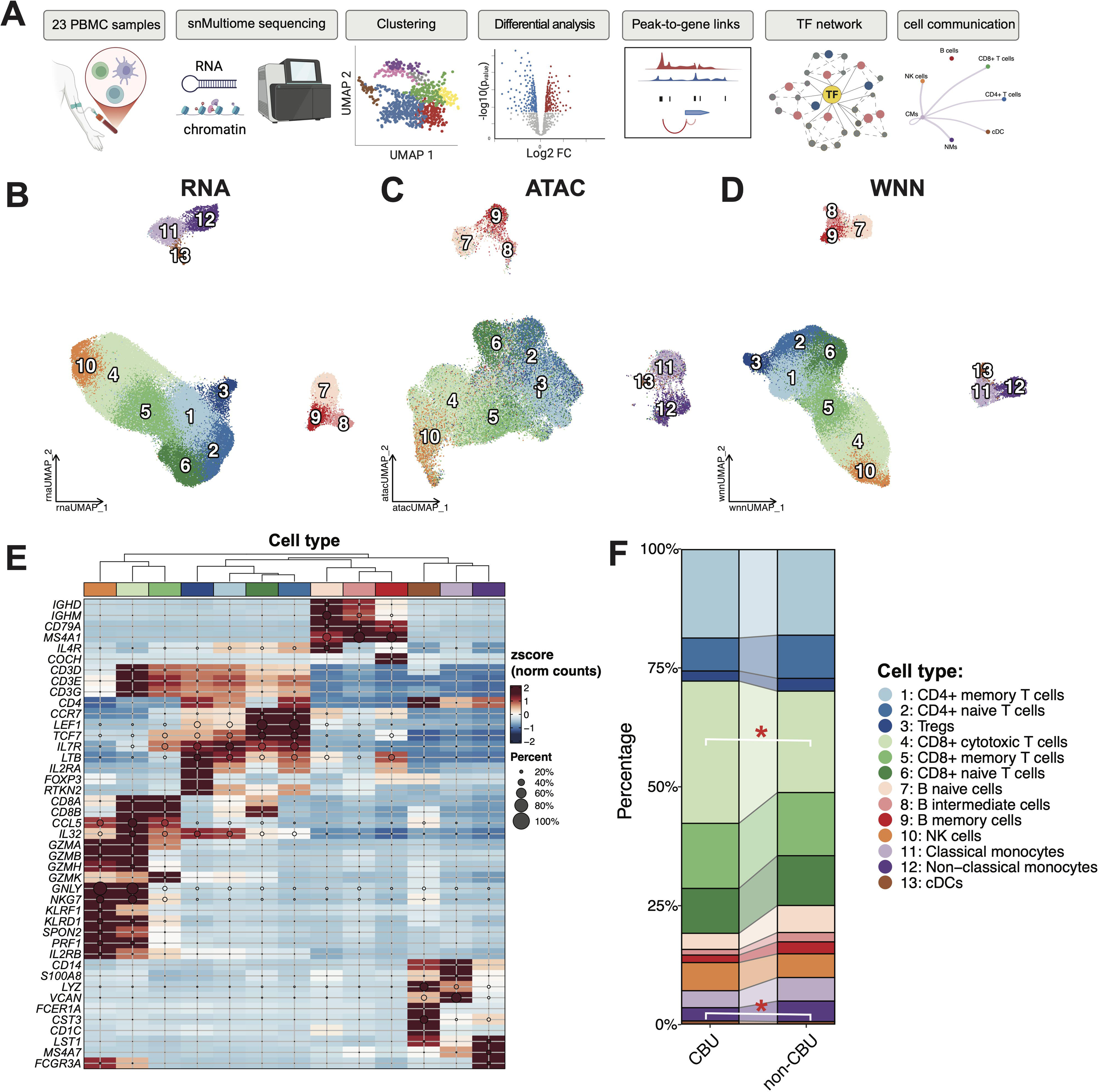
Single-nucleus (sn) multiome analysis for chronic cannabis use among people with HIV (PWH). (A) Schematic overview of the study design. FC, fold change; TF: transcription factor. (B) UMAP on snRNA-seq data, (C) UMAP on snATAC-seq data, and (D) Integration UMAP of snRNA-seq and snATAC-seq. wnn: weighted-nearest neighbor. (E) Cell type identification based on marker gene expression. Rows represent marker genes, and columns represent cell subtypes. Cell populations are clustered based on similarities of gene expression profiles. (F) Estimated cell type proportion for cannabis users (CBUs) and non-users (non-CBUs). Red asterisks mark indicates significant differences in cell type proportions between CBUs and non-CBUs (p-value< 0.05). Tregs: regulatory T cells; NK cells, natural killer cells; cDCs, conventional dendritic cells.

## Results

### Landscapes of single-nucleus transcriptomes and chromatin accessibility in peripheral blood mononuclear cells

PBMCs from PWH were profiled using a multiome approach combining snRNA-seq and snATAC-seq, including 11 CBUs and 12 non-CBUs. After rigorous quality control, a total of 23,091 genes and 172,400 peaks across 80,480 nuclei from 22 samples (1 sample was removed due to its low quality) remained for subsequent analysis **(Supplementary Table 1)**. These nuclei were clustered into 13 groups according to their transcriptome and chromatin accessibility data, along with weighted-nearest neighbor (WNN)-integrated multimodal profile^32^ **(Fig. 1B-D)**. Cell types were annotated based on expression of an established panel of marker genes for each cell cluster^32^. As shown in **Fig. 1E**, the marker genes are highly expressed in their corresponding cell types and subtypes. For example, *IGHD* and *IGHM* were highly expressed in naïve B cells, followed by intermediate B cells and memory B cells, but little expression in other cell types. The highest expression of *COCH* was in memory B cells. While *CD4* was primarily expressed in CD4+ T cells, *IL2RA* and *FOXP3* were specifically expressed on regulatory T cells (Tregs) and *LTB* were highly expressed for memory CD4+ T cells. *CCR7, LEF1*, and *TCF7* were highly expressed in naïve CD8+ T cells and *GZMK* were highly expressed in memory CD8+ T cells. Together, we identified 13 cell types including 7 major cell types and their subtypes **(Fig. 1B-D and Supplemental Fig. 1)**, CD4+ T cells (naïve CD4+ T cells, memory CD4+ T cells, and Tregs), CD8+ T cells (naive CD8+ T cells, memory CD8+ T cells, and cytotoxic CD8+ T cells), B cells (naïve B cells, intermediate B cells, and memory B cells), NK cells, classical monocytes (CMs), non-classical monocytes (NMs), and conventional dendritic cells (cDCs).

Among major cell types, CD8+ T cells showed the highest proportion of 48.5% (39,045 nuclei), followed by CD4+ T cells (23,316 nuclei, 29.0%), B cells (6,786 nuclei, 8.4%), NK cells (4,334 nuclei, 5.4%), CMs (3,510 nuclei, 4.4%), NMs (2,931 nuclei, 3.6%), and cDCs (558 nuclei, 0.7%). We compared the difference in cell type proportion between CBUs and non-CBUs based on the number of nuclei in PBMCs retained across modalities using the propeller function in the speckle R package^33^. CBUs showed an increase in cytotoxic CD8+ T cells (delta=6.73%; p=4.5e-02) and a slight decrease in NMs (delta=−2.35%; p=0.03) compared to non-CBUs **(Fig. 1F)**, suggesting that cannabis use potentially alters immune defense and the inflammatory response. There were no significant differences in the composition of other cell types between CBUs and non-CBUs.

### Differentially expressed genes between cannabis use and non-use in each cell type

We applied the MAST model^34^ implemented in Seurat^35^ to perform cell-type-specific differential transcriptomic analysis for cannabis use, adjusting for known confounding factors, with the significance threshold set at FDR < 0.05 and |avg_log2FC| > 0.15. A total of 1,234 unique DEGs were detected across 7 major cell types, with the largest number of DEGs in B cells **(Supplementary Table 2 and Fig. 2A)**. Approximately 50% (n=584) of the DEGs were cell-type-specific, indicating a high level of cell type specificity of gene expression associated with cannabis use. The remaining 650 DEGs were common in two or more cell types, while only two DEGs (<0.2%), *ARHGAP15* and *MTRNR2L12*, were in common across all seven cell types. B cells and CMs showed a greater number of unique DEGs than any other cell types, while CD4+ T cells and CD8+ T cells shared 75 DEGs, more than the number of overlapping DEGs between any other cell types **(Fig. 2A)**.

**Figure 2.**
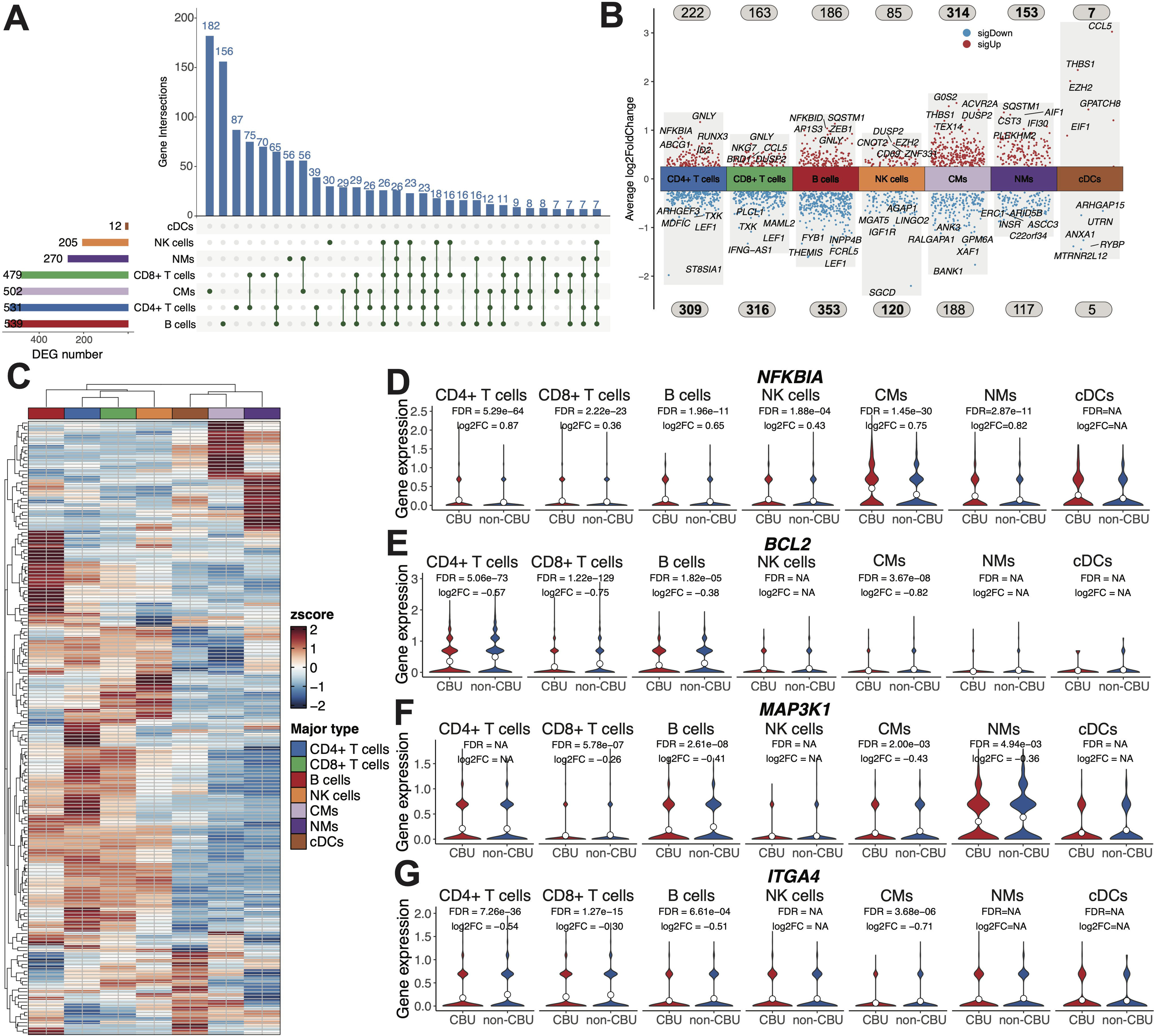
Cell-type-specific and cell-type-shared differentially expressed genes (DEGs) between cannabis users (CBUs) and non-users (non-CBUs). (A) An upset plot showing number of shared and cell-type-specific DEGs. The left bar plot shows the number of DEGs in each cell type. The top bar plot represents the number of unique or shared DEGs among cell types. Only top 30 bars are shown in the intersection plot. NK cells, natural killer cells; CMs, classical monocytes; NMs: non-classical monocytes; cDCs, conventional dendritic cells. (B) A volcano plot showing up- and down-regulated genes in each cell type. Each point corresponds to a gene, with red representing upregulated genes and blue representing downregulated genes. The gene names of top 5 up- and down-regulated DEGs in each cell type are shown. Numbers above and below the volcano plot represent the number of up- and down-regulated genes in each cell type, respectively. (C) A heatmap of gene expression showing the shared DEGs in 4 or more cell types. (D-G) Violin plots showing expression of the shared DEGs within each cell type: *NFKBIA*; *BCL2*; *MAP3K1*; and *ITGA4*. For genes that are not differentially expressed in a given cell type, the FDR and log2FC are recorded as NA. FDR, false discovery rate; FC, fold change.

Overall, we observed 1,130 genes upregulated and 1,408 genes downregulated across cell types, including genes expressed in opposite directions among cell types **(Fig. 2B)**. Lymphocytes showed more downregulated than upregulated genes between CBUs and non-CBUs, while monocytes showed the opposite pattern. **Fig. 2C** presents a heatmap of the expression level of 179 DEGs that were common among four or more cell types. Most common DEGs displayed contrasting expression patterns between lymphoid and myeloid cells, while only a few exhibited similar trends with different expression intensities. These results suggest that the effects of cannabis on the common gene expression differ between cell lineages. These common DEGs are involved in inflammatory response (e.g., *NFKBIA, NFKB1*), apoptosis (e.g., *BCL2, FAF1*), signal transduction (e.g., *MAP3K1, MAP4K4, CAMK2D*), and cell adhesion (e.g. *ITGA4, EPB41*) **(Fig. 2D-G and Supplemental Fig. 2)**. Of note, *CNR2* exhibited the highest expression in B cells **(Supplemental Fig. 3A)** and showed no differential expression between CBUs and non-CBUs in B cells but showed nominal significance in CD4+ T cells and NK cells **(Supplemental Fig. 3D)**. Two potential cannabis receptors, *GRP18* and *GRP55*, were highly expressed across cell types **(Supplemental Fig. 3B, C)**. Expression of *GPR18* showed nominal significance for cannabis exposure in CD4+ T cells and CMs **(Supplemental Fig. 3E)**, while *GPR55* was differentially expressed in CD4+ and CD8+ T cells **(Supplemental Fig. 3F)**.

### Differential expression of inflammation genes by cannabis use in each cell type

To better understand cannabis use’s immunomodulation effect in each cell type, we highlighted 36 genes involved in immune response and inflammatory function. As shown in **Fig. 3A** and **Supplementary Table 3**, these immune responsive genes exhibited distinct cell-type-specific expression patterns, which enabled discrimination of expression profiles between CBUs and non-CBUs as well as lymphocytes and monocytes. The pattern of common pro- and anti-inflammatory DEGs among cell types was complex. For example, expressions of both *NFKBIA* and *NFKB1* were increased among six out of seven cell types except cDCs (**Fig. 2D and Supplementary Fig. 2A**), suggesting that cannabis may be involved in balancing the immune response between immune inhibition (upregulated *NFKBIA*) and activation (upregulated *NFKB1*).

**Figure 3.**
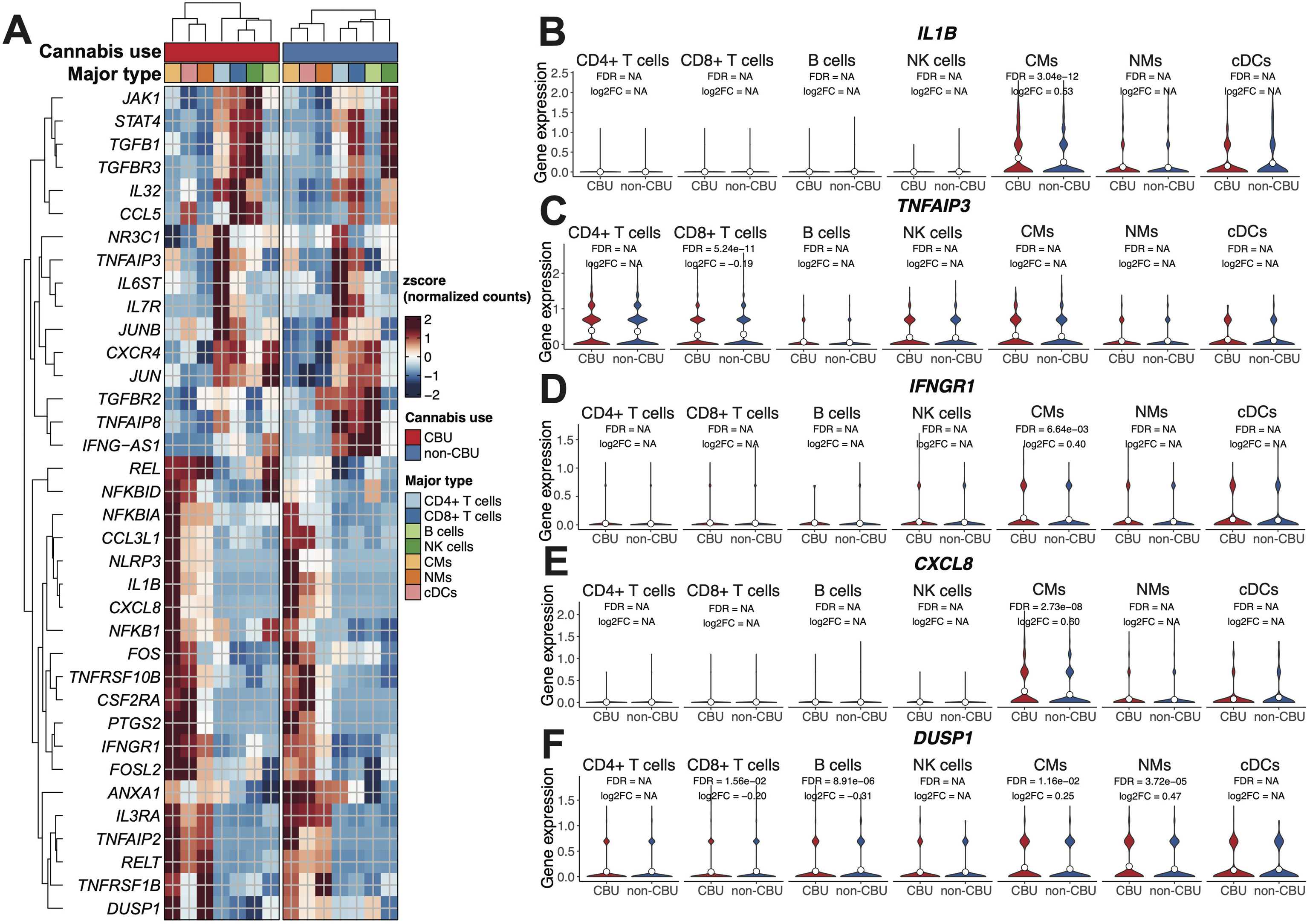
Cell-type level expression of inflammatory genes in cannabis users (CBUs) and non-users (non-CBUs). (A) A heatmap showing that distinct expression profiles of 36 inflammatory genes between CBUs and non-CBUs, as well as between lymphocytes and monocytes. NK cells, natural killer cells; CMs, classical monocytes; NMs: non-classical monocytes; cDCs, conventional dendritic cells. (B-F) Violin plots showing inflammation gene expression in CBU and non-CBU group: *IL1B*; *TNFAIP3*; *IFNGR1*; *CXCL8*; *DUSP1*. For genes that are not differentially expressed in a given cell type, the FDR and log2FC are recorded as NA. FDR, false discovery rate; FC, fold change.

A number of inflammatory DEGs were unique to cell types. Among interleukin-related genes, *IL1B* **(Fig. 3B)** and *IL3RA* were primarily differentially expressed in CMs, whereas expression of *IL32* and *IL7R* were significantly different in both CD4⁺ and CD8⁺ T cells, and *IL6ST* showed decreased expression in CD8⁺ T cells. For tumor necrosis factor (TNF) family members, *TNFAIP3* was significant in CD8⁺ T cells **(Fig. 3C)**, while *TNFRSF1B* was significant in NK cells and CMs, and both *TNFAIP2* and *TNFRSF10B* were identified in CMs. *TNFAIP8* was downregulated in lymphocytes but showed no significant difference in myeloid cells. In interferon-related genes, *IFNGR1* was significant in CMs **(Fig. 3D)**, while *IFNG-AS1* was downregulated in B cells and CD8⁺ T cells. Among transforming growth factor (TGF)-related genes, *TGFB1* was enhanced in NK cells **(Supplementary Fig. 4A)**, *TGFBR2* was found in B cells, CD4⁺ T cells, and CD8⁺ T cells, and *TGFBR3* showed enhanced expression in both CD4⁺ and CD8⁺ T cells.

Several genes encoding chemokines were differentially expressed between CBUs and non-CBUs across cell types. *CXCL8* **(Fig. 3E)** and *CCL3L1* were upregulated in CM types, but not in lymphocytes. *CCL5* was upregulated in CD8+ T cells, NK cells and cDCs **(Supplementary Fig. 4B)**. *CXCR4* was upregulated in CD4⁺ T cells, CD8⁺ T cells, and NK cells, not in monocytes **(Supplementary Fig. 4C)**. Additionally, other inflammation DEGs, like *NR3C1*, was upregulated in CD4+ T cells but downregulated in CMs **(Supplementary Fig. 4D)**. *DSUP1* is significantly downregulated by CBUs in B cells and CD8+ T cells, while it was upregulated in NMs and CMs **(Fig. 3F)**. These results indicate that long-term cannabis use is associated with widespread or cell-type-specific transcriptomic changes involved in immune and inflammatory regulation. The specific changes associated with cannabis on immune modulation likely depends on cell lineage and cell function, suggesting that comprehensive interpretation of cannabis’s pro- or anti-inflammatory effects must consider the specific cellular context.

Notably, a group of genes that were involved in HIV pathogenesis were also among the significant DEGs for cannabis use. Besides *CXCR4*, a co-receptor of HIV-1 entry in CD4+ T cells, expression of *HLA-B* was increased among CBUs compared to non-CBUs in CD4+ T cells, CD8+ T cells, B cells, and NK cells. In contrast, *RUNX1* expression was significantly downregulated in CD4+ T cells, CD8+ T cells, B cells, and NMs, but upregulated in CMs. Additionally, *ABCA1*, a host factor known to interact with the HIV-1 accessory protein Nef^36^, exhibited significantly altered expression in CMs. The HIV-1 restriction factor *SAMHD1*^37^ was identified as a DEG in CD4⁺ T cells, CD8⁺ T cells, and NK cells **(Supplementary Fig. 4E)**. The impact of cannabis uses on these genes’ expression suggest that cannabis use contributes to changes of HIV pathogenesis.

### Gene Ontology enrichment analysis in each cell type

We performed gene enrichment analysis for all DEGs, upregulated genes, and downregulated genes separately in each cell type. In the set of all DEGs, significant enrichment was observed in the pathways related to immune and inflammatory response, signal transduction, cytokine production, apoptosis, cell adhesion and activation, differentiation, proliferation (FDR<0.05**, Supplementary Table 4. a-f)**. We highlighted three categories of GO terms: immune response, cell activation and proliferation, and cell-cell adhesion **(Fig. 4)**. In the category of immune response, the pattern of the significant pathways across cell types were similar with the exception of pathways in NK cells. In the cell activation and proliferation, common pathways were observed in B cells, CD4+ T cells, and CD8+ T cells; pathways in cell-cell adherence were highly consistent across cell types.

**Figure 4.**
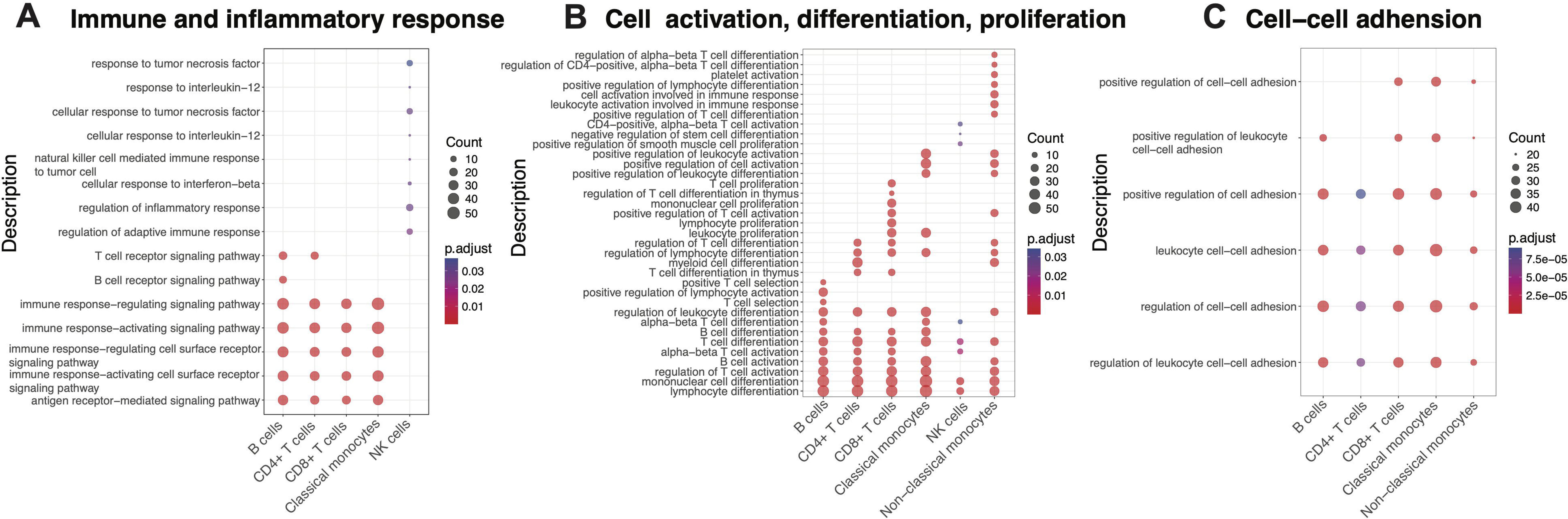
Gene Ontology (GO) analysis of Differential Expressed Genes (DEGs) between cannabis users and non-users in each cell type. Top 30 significant biological processes in each cell type are displayed (Benjamini-Hochberg adjusted p-value < 0.05). GO terms in each cell type indicate the involvement in (A) immune and inflammatory responses; (B) processes related to cell activation, differentiation, and proliferation; (C) cell-cell adhesion. Dot size corresponds to the number of genes in each GO term, and dot color indicates Benjamini-Hochberg adjusted p-value. NK cells, natural killer cells.

### Differential chromatin accessibility associated with chronic cannabis exposure in distinct cell types

We compared the chromatin accessibility landscapes between CBUs and non-CBUs by applying the LR test in Signac^38^, setting FDR < 0.05 and an absolute avg_log2 fold change greater than 0.15. A total of 4,462 unique ATAC peaks were significantly different between CBUs and non-CBUs **(Supplementary Table 5)**. The largest number of DARs were identified in CD4+ T cells, followed by CD8+ T cells and CMs **(Fig. 5A)**. No DARs were significant in cDCs, likely due to the limited cell number. Among the DARs, 3,512 peaks (78.71%) exhibited cell-type-specificity, while 950 (21.29%) were shared across multiple cell types. The largest overlapping DARs were observed between CD4+ and CD8+ T cells, consistent with their similarities in gene expression **(Fig. 2A)**. DARs were annotated and largely enriched in the promoter regions across all cell types **(Fig. 5B-C, Supplementary Table 5)**.

**Figure 5.**
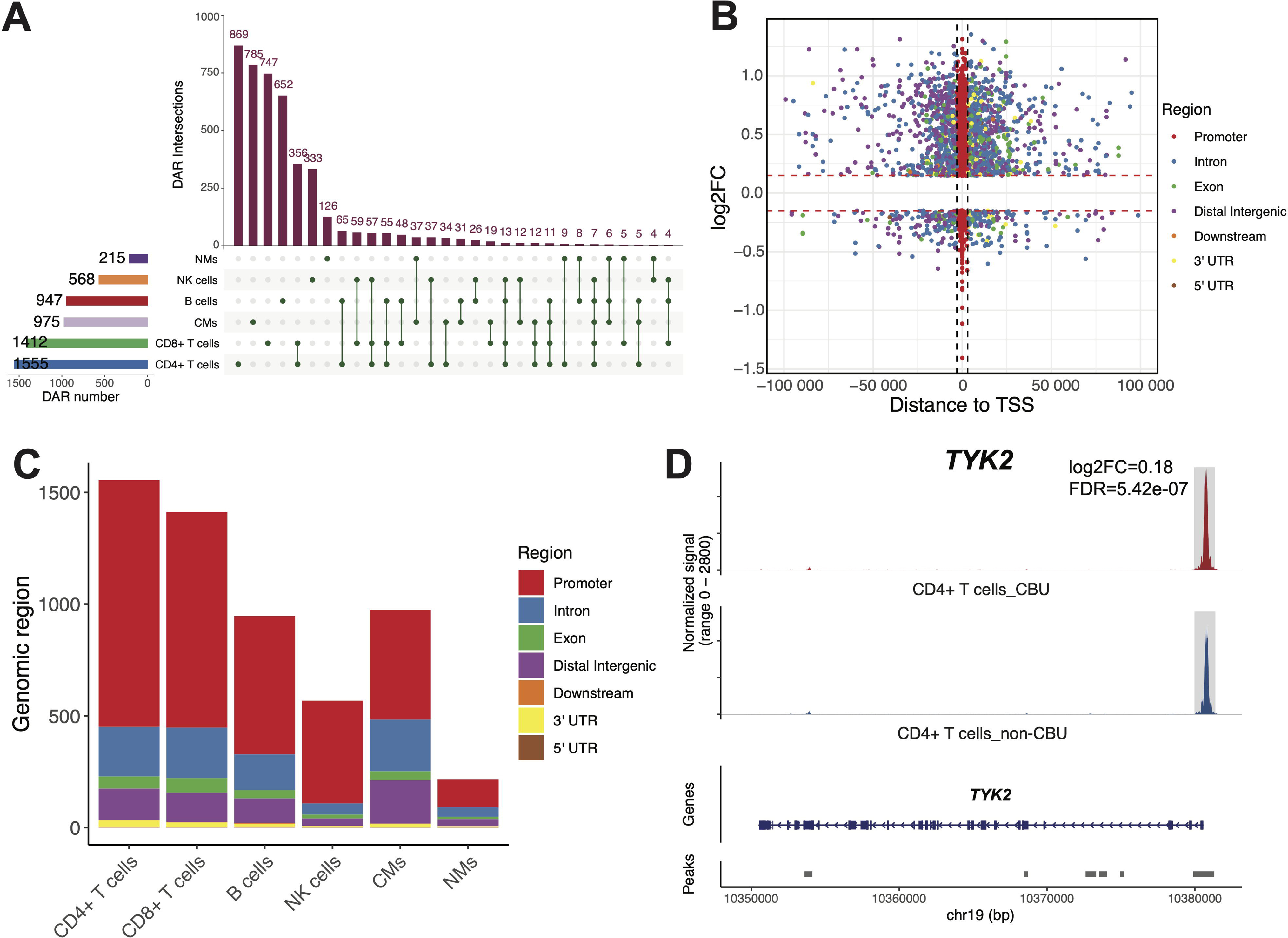
Cell-type-specific Differential Accessibility Regions (DARs) related to cannabis exposure. (A) An upset plot showing cell-type-specific and shared DARs. The bars on the left indicate the total number of DARs identified in each cell type, while the bars on the top represent the number of cell-type-specific DARs or DARs shared among different cell types. Intersection bars are ordered by frequency, and only the top 30 intersections are displayed. NK cells, natural killer cells; CMs, classical monocytes; NMs: non-classical monocytes; cDCs, conventional dendritic cells. (B) A scatter plot of DARs. Each dot represents a DAR, with colors indicating different genomic regions. The x-axis denotes the distance of each DAR from the TSS, and the y-axis represents the log2 fold change (FC). The promoter region is defined as ±3 kb around the TSS. TSS, transcription start site; UTR, untranslated region. (C) A bar plot showing the distribution of DARs across different genomic regions in each cell type. Colors indicate different genomic regions. (D) An example of a DAR located in the promoter of its nearest gene *TYK2* in CD4+ T cells. The top panel shows chromatin accessibility, with cannabis user (CBU) shown in red and non-user (non-CBU) in blue. The bottom panel displays the gene coordinates and peak annotation. DAR is highlighted with gray background. FDR, false discovery rate.

Multiple DAR-proximal genes were involved in immune response and inflammation regulation in different cell types. In CD4+ T cells, a DAR near *TYK2* from the Janus kinase (JAK) gene family showed an increased chromatin accessibility in CD4+ T cells in CBUs compared to non-CBUs **(Fig. 5D).** *TNFRSF1A* is crucial for regulating inflammation and activating immune cells by binding to tumor necrosis factor alpha (TNF-α) ^39^. A DAR which is tightly linked to *TNFRSF1A* exhibited increased chromatin accessibility in CBUs **(Supplementary Fig. 5B)**. Additional DAR-proximal genes in CD4+ T cells included *TNFRSF1B*, *TNFSF8*, *TNFRSF12A*, *TNFRSF18* and *FOSL1* **(Supplementary Fig. 5C)**, *FOSL2*, *IL23A*, *CXCR5*, *PDE4A*, *PDE4B* **(Supplementary Table 5),** which all play critical roles in inflammation. In CMs, a subset of DARs positioned near immune and inflammatory genes included *NFKBIA*, *NLRP3*, *PDE4A*, *PDE4B*, *MAP3K14*, *TNIP3*, *CCL20*, *IL1A*, *IL18*, *IL12RB1*, *IL6R*, *IL3RA*, *IL10RA*, and *ANXA1.* In particular, a more accessible DAR was observed near the *NLRP3* gene, accompanied by DARs with enhanced accessibility overlapped with *IL10RA* and *PDE4A* **(Supplementary Fig. 5D-F)**. These findings indicated a potential link between chromatin accessibility changes and the regulation of immune- and inflammation-related genes influenced by cannabis use.

### Cell type-specific *cis*-regulation altered by chronic cannabis use

To explore the regulatory mechanisms linking gene expression to chromatin architecture, we examined the candidate *cis*-regulatory elements (cCREs) of DEGs by estimating Pearson correlation between expression level of DEGs and chromatin accessibility of DARs in each cell type. A total of 242 unique peak-to-gene (p2g) links between DEGs and DARs were identified across six major cell types (**Supplementary Table 6**). The regulatory role of the most CRE-gene links (N=230, 95.04%) showed cell type specificity, while small subset (N=12, 4.96%) was shared in more than one cell types (**Supplementary Fig. 6A**).

We found that cCREs played a role in cannabis-associated immune and inflammatory gene expression. An increased expression of *NFKBIA* in CBUs was positively correlated with the chromatin accessibility of its potential enhancer region (DAR: chr14-35415732-35417260, r=0.08, FDR=1.96e-08, **Fig. 6A)** in CD4+ T cells. Similarly, the expression of *NFKB1* was significantly associated with chromatin accessibility of a DAR (chr4-102500305-102502522) at its promoter (r=0.07, FDR=9.03e-05, **Supplementary Table 6)** in CD4+ T cells. Another significant link in CD4+ T cells was a DAR near *GATA3*, a TF crucial for the development and function of immune cells, particularly T helper 2 (Th2) cells. DAR: chr10-8415849-8417160 within the putative distal regulatory region of *GATA3* showed increased accessibility in CBUs, accompanied by elevated *GATA3* expression in CD4+ T cells **(Fig. 6B)**. The correlation analysis showed a significant association between the transcription level of *GATA3* and chromatin accessibility (FDR=1.71e-14, r=0.10). Moreover, the expression level of *BCL2* involved in apoptosis, the signal transduction gene *SQSTM1* and regulators of immune response such as *TCF7, HLA-B*, and *CD69*, genes play critical roles in inflammation such as *RUNX3, NR3C1* and *JUNB*, were significantly correlated to differentially accessible CRE’s chromatin activity in CD4+ T cells **(Supplementary Fig. 6B-E and Supplementary Table 6)**.

**Figure 6.**
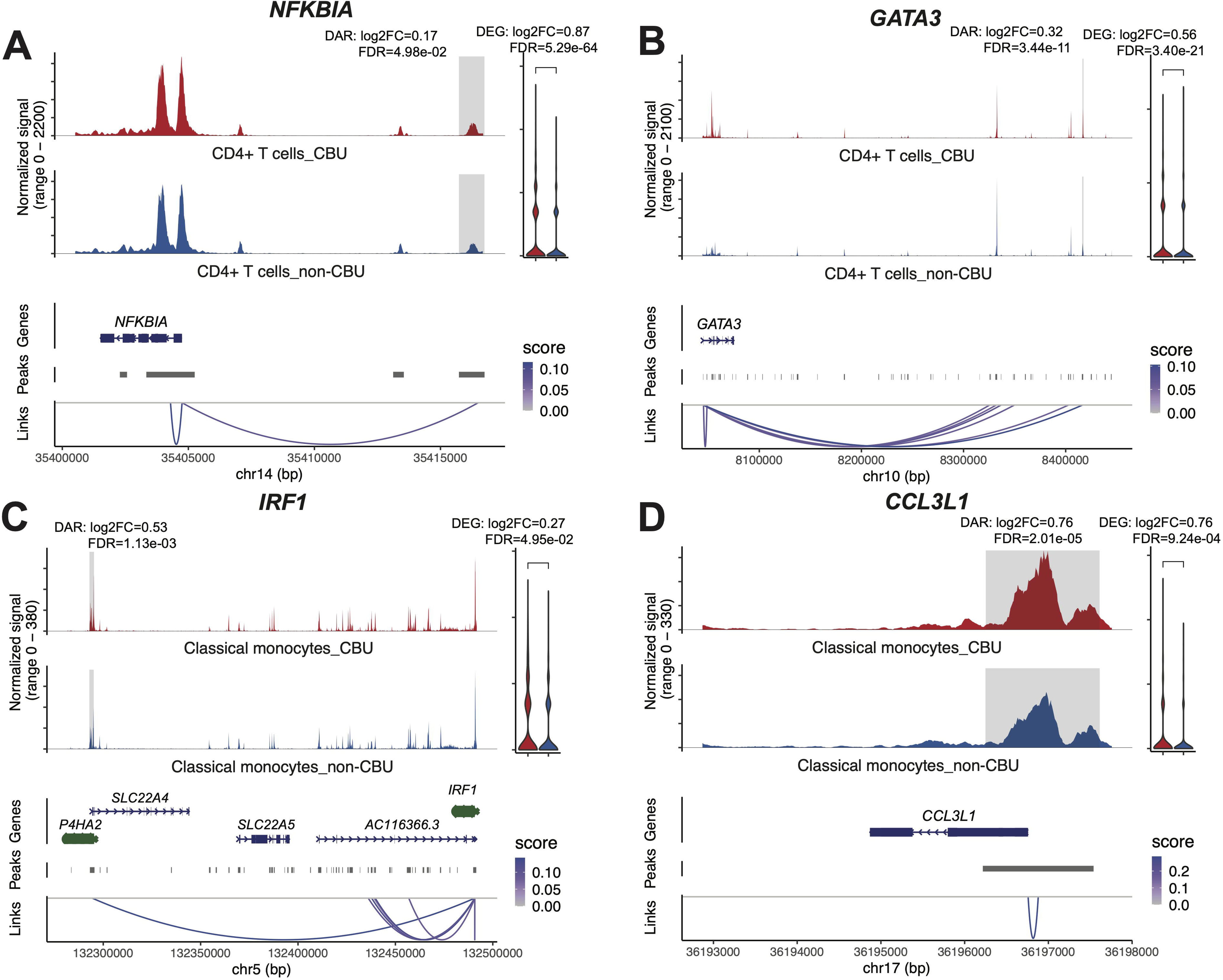
*cis*-regulatory interaction between Differential Accessibility Region (DAR) and Differential Expressed Gene (DEG) in individual cell types. Coverage plots showing the correlations between gene expression of DEGs and chromatin accessibility of DARs. Top panel: chromatin accessibility in cannabis users (CBUs, red) and non-cannabis users (non-CBUs, blue), with DARs shown in gray. Middle panel: gene and peak annotations. Bottom panel: linkage between DEGs and DARs. (A) *NFKBIA* in CD4+ T cells; (B) *GATA3* in CD4+ T cells; (C) *IRF1* in classic monocytes; and (D) *CCL3L1* in classic monocytes. FDR: false discovery rate; FC: fold change.

*IRF1*, a crucial regulator of the immune system and plays a vital role in cellular processes such as inflammation, cell growth, and apoptosis^40,41^. Expression of *IRF1* was upregulated in CBUs compared to non-CBUs and chromatin accessibility of a putative enhancer of this gene was greater in CBUs than non-CBUs in CMs **(Fig. 6C)**. Moreover, *IRF1* expression was significantly correlated with the chromatin accessibility of this enhancer region in CM (r=0.14, FDR=9.74e-04). A chemokine gene *CCL3L1*, which inhibits HIV entry, playing a crucial role in HIV/AIDS pathogenesis and susceptibility^42,43^ was upregulated in CBUs. Correspondingly, DARs located at its promoter (chr17-36196217-36197540) was more accessible in CBUs compared to non-CBUs. Significant correlation (r=0.29, FDR=1.02e-15) was found between the chromatin openness of the DAR and the expression level of *CCL3L1* **(Fig.6D)**. In addition, the expression of NF-κB pathway-related genes, including *NFKBIA, NFKB1, NFKBIZ,* and *REL*, were closely correlated with their corresponding cCREs’ chromatin accessibility, suggesting altered epigenetic regulation of NF-κB signaling in CBUs. Similar correlations were also observed in inflammation genes *NLRP3, JUND, FOS, ATF3*, and *IL1B*, apoptosis gene *BID*, HIV-related genes *NR4A1* and *NFAT5* (**Supplementary Fig. 6F-H**). Together, these results suggest a potential link between chromatin and gene expression regulation in immune response, inflammatory response, apoptosis, and HIV susceptibility.

### Critical transcription factors regulate differentially expressed genes for cannabis use

To provide a more comprehensive view of gene expression regulation beyond cCREs, we assessed the influence of chronic cannabis use on transcription factor binding motif (TFBM) activity and the co-expression network. Differentially active TFBMs corresponding to cannabis exposure were determined within each cell type **(Supplementary Table 7)**. Comparative analysis revealed that both the transcriptional expression and motif activity of several immune-regulatory TFs, differed significantly between CBUs and non-CBUs. In CD4+ T cells, TFBMs of *NFKB1* and *JUNB* were more active and their RNA abondance were higher in CBUs, while *TCF7* exhibited decreased TF motif activity and gene expression in CBUs **(Fig. 7A-F)**. TFBMs of *JUND* and *FOS* were more active in CBUs in CMs **(Fig. 7G-H)**. Correspondingly, the expression levels of *JUND* and *FOS* were increased in CBUs **(Fig. 7J-K)**. *NR4A1* motifs showed decreased activity but elevated RNA abundance in CBUs **(Fig. 7I, L)**, suggesting *NR4A1* could function as a transcription suppressor in PWH. These findings suggest potential shifts in transcription factor binding may underline functional differences of immune response between CBUs and non-CBUs.

**Figure 7.**
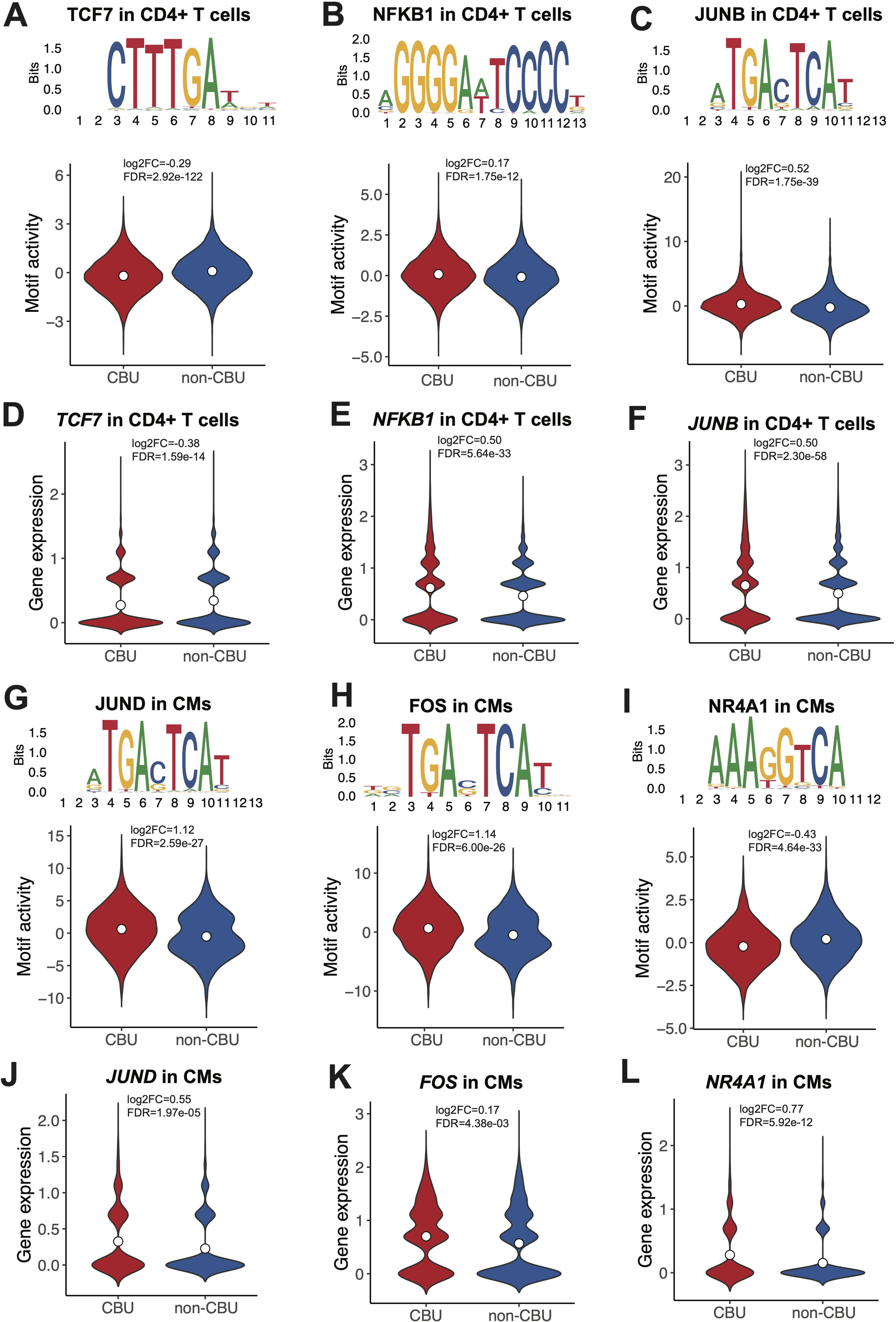
Differentially active transcription factor binding motifs and their corresponding gene expression between cannabis users (CBUs) and non-users (non-CBUs) in CD4+ T cells and classical monocytes. **(A-F).** Motif activity and gene expression in CD4+ T cells: TCF7; NFKB1; and JUNB. (G-L) motif activity and gene expression in classic monocytes: JUND; FOS; and NR4A1. TF binding motif shown as motif logo above. CMs: classical monocytes. FDR: false discovery rate; FC: fold change.

We developed TF regulatory networks for the aforementioned TFBMs that exhibited differential activity between CBUs and non-CBUs, tailored to individual cell type. We defined the target genes of a specific TF as those with accessible binding motifs for the TF in their cCREs, and whose expression is significantly associated with the TFBM’s activity. Within the constructed networks, we observed multiple DEGs under the influence of differentially active TFBMs in both CD4+ T cells **(Fig. 8A)** and CMs **(Fig. 8B)**. Among target genes, multiple immune and inflammation genes are observed, such as *JUND, IRF1, IL7R, RUNX3, TGFBR2, TCF12, NFKBIA, FOSB*, and *JUN* in CD4+ T cells and *IL1B, NFKB1, NFKBIA, CXCL8, IL3RA, NLRP3*, and *ANXA1* in CMs. Our data implied that the dysregulation of gene expression in CBUs may result from altered TFBM activity and chronic cannabis use could influence the regulatory network of immune and inflammatory response.

**Figure 8.**
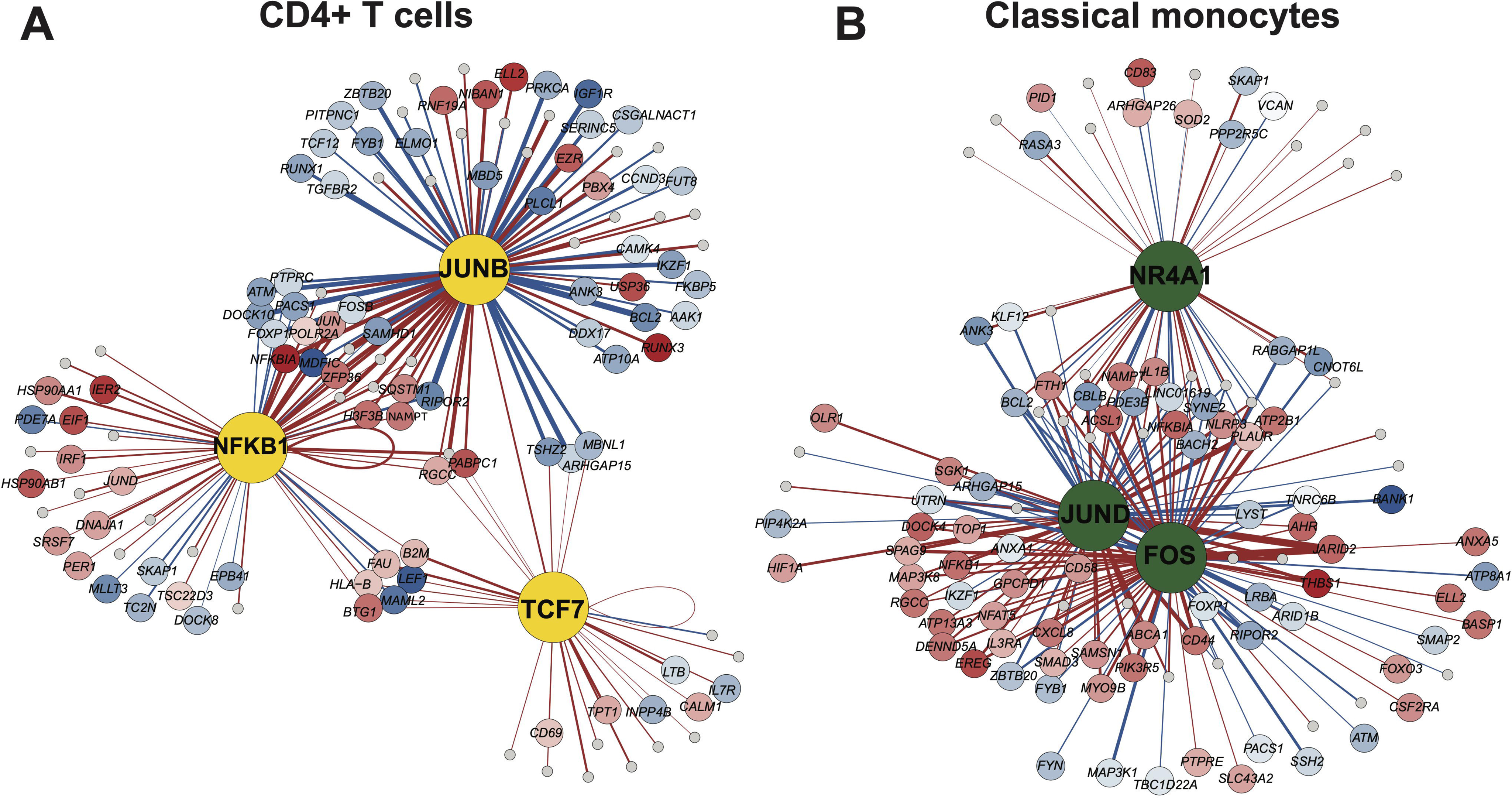
Transcription factor (TF) regulatory network in CD4+ T cells and classical monocytes. The largest dot in each network represents transcription factor. Dots indicate upregulated (red) and downregulated (blue) genes between cannabis users (CBUs) and non-users (non-CBUs), with color intensity reflecting the magnitude of the absolute log2 fold change. Gray dots represent TF target genes that are not differentially expressed between CBUs and non-CBUs. The thickness of the edges between the TF and its target genes represents the correlation between transcription factor binding motif activity and its target gene expression. Red edges indicate positive correlation, while blue edges indicate negative correlation. (A) TF-gene network in CD4+ T cells; (B)TF-gene network in classical monocytes.

### Cannabis use altered cell-cell communications between lymphoid and myeloid cell lineages

Cell-cell communication networks were presented in **Fig. 9A and Supplementary Fig. 7A-C**. CD8+ T cells showed the most changes in incoming signals, followed by B cells, while CMs had the most altered outgoing signals, followed by NMs (**Fig. 9B and Supplementary Fig. 7D**). When CMs functioned as signal senders, their interaction strength increased with most immune cell types, except cDCs. In contrast, as signal receivers, CMs exhibited enhanced interaction strength with all other cell populations. Similarly, when CD8⁺ T cells act as signal sources, showed universally increased interaction strength with other immune cell types except NK cells. Notably, NK cells, when receiving signals, demonstrated reduced interaction strength with lymphoid cells, but increased interaction strength with myeloid cells **(Fig. 9B)**.

**Figure 9.**
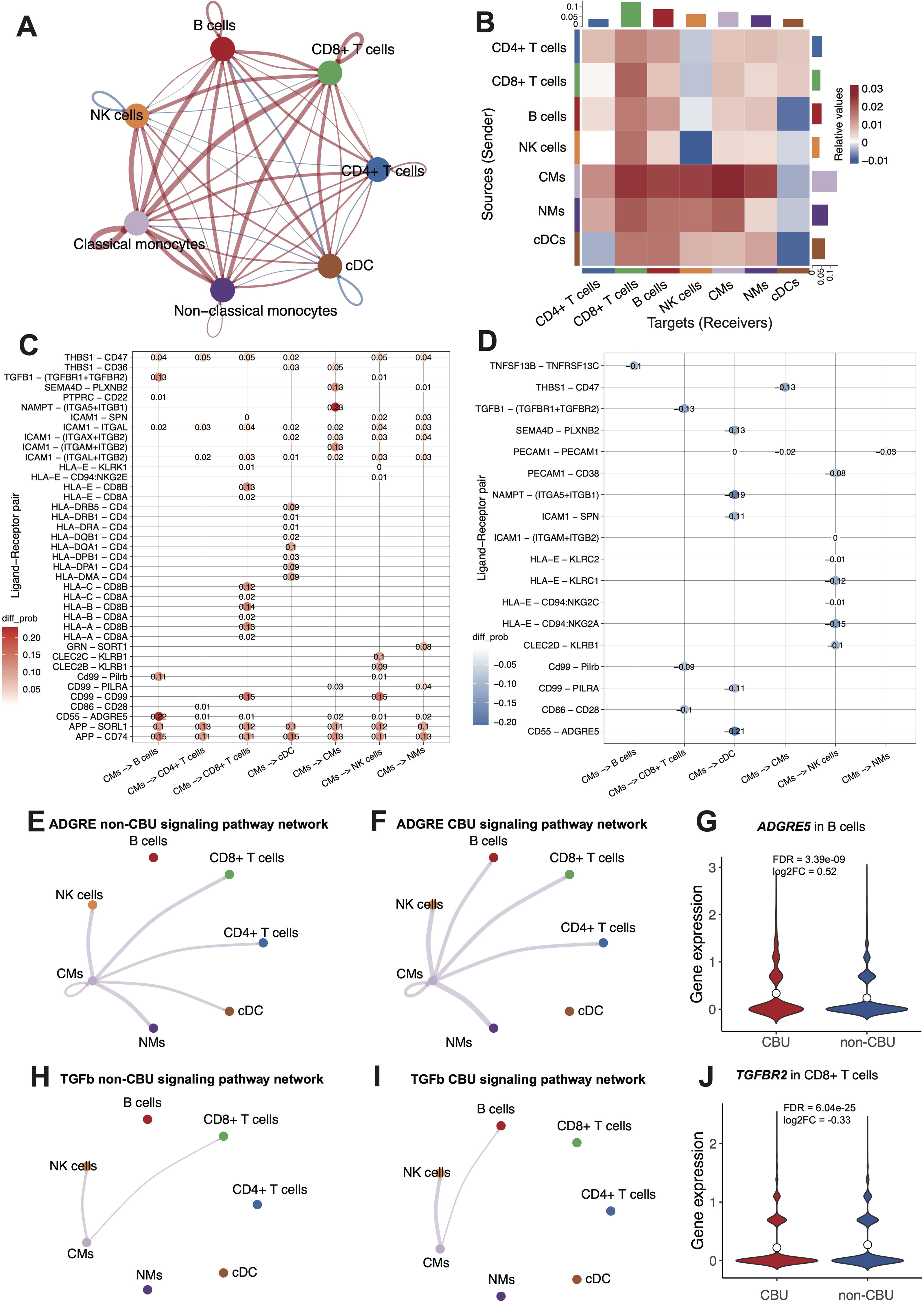
Cell-cell communications among cell types. (A) A circle plot representing the overall interaction network between different cell types, in which edge represents increased (red) or decreased (blue) signaling and the thickness of the edge line indicates a strength of the signaling; (B) A heatmap showing the different interaction strength between “sender” cell type (y-axis) and receiver cell type (x-axis). The colored bar plots represent a total incoming signaling for each column (top) and a total outgoing signaling for each row (right). Color shades indicate increased (red) or decreased (blue) values of interaction in cannabis users (CBUs) relative to non-users (non-CBUs). NK cells, natural killer cells; CMs, classical monocytes; NMs: non-classical monocytes; cDCs, conventional dendritic cells. (C) and (D): Dot plots detailing specific ligand-receptor pairs and their communication probability changes across various cell-cell interaction pairs. (E) and (F) Hierarchy plots demonstrating ADGRE signaling pathway network of (E) non-CBUs and (F) CBUs; (G) A violin plot showing expression level of *ADGRE5* in B cells between CBUs and non-CBUs; (H) and (I) Hierarchy plots showing TGFb signaling pathway network for (H) non-CBUs and for (I) CBUs; and (J) A violin plot showing expression level of *TGFBR2* in CD8+ T cells between CBUs and non-CBUs. FDR: false discovery rate; FC: fold change.

When CMs acted as signal sender, we identified multiple ligand-receptor pairs with increased **(Fig. 9C)** or decreased **(Fig. 9D)** interaction strength between CMs and other cell types. Notably, the ADGRE signaling pathway between CMs and B cells was enhanced in the CBU group (*CD55 - ADGRE5*, **Fig. 9C, E, F)**, which may be attributed to the elevated expression of *ADGRE5* in B cells **(Fig. 9G)**. *ADGRE5* (also known as *CD97*) is an adhesion G protein-coupled receptor involved in inflammation by regulating immune cell adhesion, migration, and activation. In contrast, the TGF-β signaling pathway (*TGFB1-TGFBR1+TGFBR2*) between CMs and CD8⁺ T cells was reduced in the CBU group **(Fig. 9D, H, I)**, potentially driven by the decreased expression of *TGFBR2* in CD8⁺ T cells **(Fig. 9J)**.

## Discussion

This study integrates snRNA-seq and snATAC-seq profiles to comprehensively evaluate the impacts of chronic cannabis exposure on gene expression and chromatin accessibility in immune cells from PWH. Overcoming the limitations of previous studies in bulk cells, we found significant differences between CBUs and non-CBUs in gene expression, chromatin accessibility, TFBM activity as well as cell-cell communication at single cell level. We identified hundreds of cannabis use-associated DEGs that were regulated by changes in chromatin accessibility and inferred activity of putative TFBMs in major cell types. Cannabis’ immunomodulation effect in gene expression is highly context-dependent across cell lineages and cell types, particularly genes involving in inflammation. We observed evidence of increased cell communication among CD8+ T cells, B cells, and monocytes, and each in turn with other cell types among cannabis users. Together, the converging evidence from multi-layer analyses underscores the dynamic and complex effects of cannabis on immune system in the HIV-infected host genome, showing suppressed immune function in some cell types while proinflammation was observed in others. Understanding chronic cannabis uses nuanced but presumably persistent immunomodulatory effect may have profound clinical implications. To the best of our knowledge, this is the first study applying sn-multiomic technology to cannabis use.

Approximately 50% of the DEGs were specific to individual cell types. Similarly, most DARs (3,512 peaks, 78.71%) and peak-to-gene links (230, 93.36%) are specific to particular cell types. The TF regulatory networks involve distinct differentially accessible TFBMs and their corresponding target genes are also unique to different cell types. Thus, gene expression patterns, chromatin states, and both *cis*- and *trans*-effects of chromatin accessibility on gene regulation collectively underscore the pronounced cell-type specificity and exhibit considerable heterogeneity. The detailed multi-level characterization of transcriptomes and chromatin landscapes within each cell type provides a precise profile of cannabis’s broad effects on immune cells, which cannot be achieved through bulk transcriptome analyses.

At the transcriptome level, consistent with our previous report on a single cell transcriptome analysis of acute THC exposure^31^, we identified a large number of DEGs associated with chronic cannabis use in cell-type-specific fashion. Many of these DEGs were implicated in immune and inflammatory responses, cytokine production, apoptosis, and signal transduction-pathways previously implicated in cannabis-related studies^44–46^. Interestingly, these effects on gene expression exhibit a variety of distinct gene regulation profiles between lymphocytes and monocytes, particularly genes involved in immune modulation. Cannabis appears to have a dual role with both pro- and anti-inflammatory effects across cell types with a considerable amount of cell type-specific effects. Cannabis use upregulated inflammatory genes *NFKB1* in CD4+ T cells, chemokine genes *CCL5* in CD8+ T cells, NK cells and cDCs*, CCL3L1*, and *CXCL8* in CMs, as well as TNF family genes including *TNFRSF1B* and *TNFRSF10B* in CMs. These findings suggest that cannabis use could directly increase inflammatory gene expression, enhance chemokine-mediate immune cell activation and recruitment, and increase inflammatory function via upregulation of inflammation gene expression. On the other hand, cannabis use upregulated genes involving negative immune regulation (*NFKBIA* and *ZFP36* in CD4+ T cells, NK cells and NMs), upregulated anti-inflammatory genes related to type I and type II interferon responses in B cells and CMs, and downregulated pro-inflammatory cytokine pathways, such as interleukin 2 production in certain cell types (leading to increased anti-inflammatory cytokine production). These findings are consistent with evidence reported in the literature regarding cannabis exposure’s anti-inflammatory effects. The dual pro-and anti-inflammatory effects are highly cell-type-specific. This complex interplay suggests that while cannabis maybe associated with suppressed immune reaction and decreased inflammation in certain immune cell types, it may be linked to promoted inflammation process in other cell types. The net effect of cannabis on inflammation may be determined by the balance of its action across different cell types and specific pathways involved. We speculate that the disrupted coordination of pro- and anti-inflammatory processes, while showing a net anti-inflammatory profile, may in fact lead to unfavorable immune functioning in the long term such as in the setting of chronic cannabis use. Our findings highlighted the need for more targeted and nuanced research to fully understand its immunomodulatory effects.

Chromatin accessibility analysis has revealed the cell-type-specific immune modulation that chronic cannabis use exerts is similar to its impacts on immune cell transcriptomes. We identified a substantial number of DARs associated with cannabis use and their proximity to genes related to immune response and inflammation. One example is *TYK2*, which is involved in the signaling of cytokines such as IL-12, IL-23, IFN-α/β, IL-6, and IL-10^47,48^. We found that *TYK2* showed greater accessibility in CBUs in CD4+ T cells, suggesting that cannabis could increase chromatin openness near *TYK2* that leads to increased pro-inflammatory gene expression, at least in the context of HIV infection. Other DAR-proximal genes linked to cannabis use included *TNFRSF1A, TNFRSF1B*, *TNFSF8*, *TNFRSF12A*, *TNFRSF18, FOSL1*, *FOSL2*, *IL23A*, *CXCR5*, *PDE4A*, and *PDE4B*.

Perhaps stronger evidence is from cCRE-gene correlation of DARs to DEGs in *cis*-relationships. In CD4+ T cells, we observed that *GATA3* was upregulated in CBUs. A previous report that THC increases the expression of GATA3 that shifts T helper cell balance^49^. Notably, the expression of *GATA3* was correlated with the accessibility of its putative enhancer region. Our analysis indicates that an increased *GATA3* expression is likely due to an increase chromatin openness by cannabis exposure. Furthermore, in CMs, expression of chemokine gene *CCL3L1,* which inhibits HIV entry, was significant correlated with its CREs’ chromatin accessibility. The same applies to NF-κB pathway-related genes, including *NFKBIA*, *NFKB1*, *NFKBIZ,* and *REL*, suggesting epigenetic regulation of NF-κB signaling in CBUs. In addition, *IL1B* and *IRF1* are key regulators of innate immune responses. *IL1B* encodes for a pro-inflammatory cytokine^50^, while *IRF1* is a transcription factor that orchestrates inflammatory gene expression^51^. The transcriptional activation of these genes strongly correlates with chromatin structure, suggesting that epigenetic mechanisms likely modulate gene expression during immune activation and inflammatory responses which were altered by chronic cannabis use.

Another layer of evidence regarding cannabis use’s immune regulatory effects is its influence on TF activity and their target gene expression. We identified changes in TFBM activity in the CBUs compared to the non-CBU group, and many of the TF targeted genes showed differential expression. These results suggest that the DEG alterations in CBUs appear to be at least in part the result of chromatin accessibility alterations at their cCREs and dysfunction of TFBM activity. This multi-layered regulation could suggest that TF binding is modulated by chromatin accessibility state, which impacts transcription activation. Such coordinated transcriptional and epigenetic regulations likely underline the CBU-specific expression pattern of DEGs. These results highlight the unique capability of combining both DNA accessibility and gene expression measurements from the same cell to enable comprehensive genomic insights.

In the context of chronic HIV infection, cannabis use appears to exert changes in the expression of genes involving HIV-1 entry and latency. *CXCR4*, a gene encoding for a co-receptor of HIV-1^52,53^, is upregulated by cannabis exposure in CD4+ T cells. Conversely, *SAMHD1*, a gene preventing HIV-1 infection in resting CD4+ T cells^54^, is downregulated in CD4+ T cells. Previous studies showed that *SAMHD1* negatively modulates reactivation of viral latency in CD4^+^ T cells^55^. Furthermore, the NF-κB signaling pathway, encompassing the *NFKB1* and *NFKBIA* genes whose expression were changed by chronic cannabis use, plays a pivotal yet complex role in HIV infection and the establishment and maintenance of viral latency^56,57^. These findings indicate that cannabis use leads to transcriptional alterations in HIV-related genes, reshaping immune regulation and inflammation response in HIV infection. These findings highlight the complexity of cannabis’s biological effects, indicating that its impact on immune function in PWH cannot be generalized across all cell types. Therapeutic use of cannabis or cannabinoids needs to take into account these nuanced and context-dependent effects.

We identified differential patterns of cell-cell communication between chronic cannabis users and non-users, whereby CD8+ T cells and B cells displayed increased incoming communication while monocytes displayed increased outgoing communications with other cell types. We highlighted two signaling pathways influencing the observed cell-cell interactions. Specifically, the ADGRE (CD97)-CD55 interaction increased between CMs and B cells, showing cannabis use’s role in cell adhesion, migration, and activation^58^. Enhanced activation of this pathway could lead to increased recruitment and retention of immune cells at inflammation sites, contributing to chronic inflammatory conditions. In the context of chronic HIV infection, chronic cannabis may exacerbate inflammatory responses in some individuals who already experience a sustained low-degree inflammatory status. However, we also observed the reduction of TGF-β pathway expression in cannabis users, potentially driven by decreased expression of *TGFBR2* in CD8+ T cells. TGF-β signaling is critical for immune regulation, including the suppression of inflammatory responses and maintaining immune homeostasis^59^. The findings suggest that cannabis use exhibits coordinated effects via regulating interactions of different cell types.

Despite several strengths to this study, there are also limitations warranting acknowledgement. The idea to investigate cannabis use’s effect in PWH is to understand the inflammatory effects of sustained cannabis use in the setting of chronic immune activation due to HIV infection. For this purpose, the samples went through strict selection from over 2,000 subjects to match CBUs and non-CBUs. All samples were collected from PWH taking antiretroviral therapy and with viral suppression for at least two years. We expect minimal confounding due to HIV-1 load and other medical comorbidities. However, lacking the samples from persons who are not living with HIV infection limits the generalization of cannabis’s immune modulation effects to people without HIV. All subjects were women, which precluded generalizing findings to men although minimized confounding by sex. Furthermore, cannabis contains a unique combination of up to and more than 550 components. In the current study, cannabis use was defined using self-report of use over the prior six months, which does not allow us to estimate the concentration, active components, or complex pattern of exposure of use. Additionally, the presence of polysubstance use and other unknown factors may confound the findings. Furthermore, some participants classified as current non-users were former users, which could influence the interpretation of potential epigenetic changes. These limitations warrant consideration in interpretation of the findings. Future studies in larger sample sizes to replicate the findings are warranted.

In summary, our study addresses the contentious issue of whether cannabis consistently exerts anti-inflammatory effects and whether it could benefit PWH. Our research from the multi-layered omic evidence indicates that chronic cannabis exposure is linked to changes in the transcriptome and epigenome, with the immune modulation effect varying depending on immune cell context. The findings demonstrate that cannabis use may result in immune dysfunction in the setting of HIV infection by profoundly modulating transcriptome in different immune cells and changes communication. A nuanced understanding of these effects is crucial for clinicians managing patients who use cannabis, particularly those with preexisting immune-related conditions. Further research is essential to fully elucidate these interactions to begin to understand how chronic cannabis use may impact managing chronic HIV infection along with comorbid conditions.

## Methods

### PWH PBMC samples

Samples from selected participants were obtained from the MACS/WIHS Combined Cohort Study (MWCCS). The study design, study protocol, cohort characteristics, and specimen repository were described previously^60^. Frozen viable PBMC samples (n = 23) collected from PWH who had maintained viral suppression under antiretroviral therapy for a minimum of two years were received from the MWCCS Biorepository. This PWH cohort consisted of clinically matched CBUs and non-CBUs. Eleven participants with a history of long-term (at least 6 months) cannabis use and current use at the time of sample collection were included in the CBU group. Twelve participants reported never use (n=3) or former use (ranging from 6 months to 11 years since use). Of the CBUs, 8 participants also reported concurrent cocaine use, whereas 5 individuals in the non-CBU group reported cocaine use. Cocaine use was covariate in the analyses. Demographic data are presented in **Supplementary Table 1**.

### Nuclear dissociation and library preparation

PBMCs in freezing media were thawed according to PBMCs/Cell Lines-Direct Media Thawing Protocol (10X Genomics CG000447). Quantity and viability of the cells were measured using the Luna FX7 Automated Cell Counter (Catalog #L70001, Logos Biosystems) with Acridine Orange/Propidium Iodide (AO/PI) fluorescence staining. The cells were then pelleted, resuspended in 1mL of 10% nuclei isolation reagent (NIR), 1 part NIR and 9 parts nuclei storage reagent (NSR) (Precision Cell Systems, S200 Reagent) was loaded into a chilled Nuclei Isolation Cartridge Plus (Catalog #100-289-043, Precision Cell Systems) with RNase inhibitor and isolated using the manufacturer’s recommended nuclei isolation protocol on the S200 Nuclei Isolation System (Precision Cell Systems). Nuclei quality and quantity were measured using the Luna FX7 Automated Cell Counter with (AO/PI) fluorescence staining. The nuclei were then used as input for the Chromium Single Cell Multiome ATAC + Gene Expression protocol (10X Genomics). The samples were processed and sequenced according to the manufacturer’s instructions.

### Quality control for both snRNA-seq and snATAC-seq data

Cell Ranger ARC (v2.0.2) was used to process snRNA-seq and snATAC-seq data with default parameters, using the GRCh38 reference genome. Unless otherwise specified, Seurat v5.2.0^32^ was used to process and analyze the snRNA-seq data, while Signac (v1.14.0)^38^ was employed for the analysis of snATAC-seq data. Ambient RNA was computationally estimated by R package decontX (v1.0.0)^61^. Genes annotated as mitochondrial or ribosomal were filtered out from the count matrix prior to normalization and downstream analysis. Low-quality nuclei were removed based on both snRNA-seq and snATAC-seq QC metrics (nFeature_RNA<200, nCount_RNA<300 or >95th percentile of nCount_RNA, %Mitochondrial genes> 10%, %Ribosomal genes>30%, %ambient RNA>20%, nCount_ATAC<1500 or >40,000, blacklist_ratio> 1%, TSS.enrichment<4, nucleosome_signal>1). Doublets were identified and removed using DoubletFinder R package (v2.0.4)^62^ based on snRNA-seq data. Only cells that passed quality control in both the snRNA-seq and snATAC-seq modalities were retained for downstream analyses. In addition, during cell clustering, 9 RNA clusters and 3 ATAC clusters were removed due to inconsistent clustering between RNA and ATAC modalities. Finally, a total of 80,480 nuclei were retained for further analysis.

### Single nucleus RNA data analysis workflow

After removing low quality nuclei, SCTransform() normalization with default parameters was applied to the filtered RNA count matrix. Linear dimensional reduction was performed by RunPCA() function, followed by batch effect correction using harmony v1.2.3^63^ through RunHarmony() function on the SCT assay. A shared nearest neighbor graph was constructed with the first 30 dimensions using the FindNeighbors() function and cells were clustered using the Louvain algorithm via the FindClusters() function (resolution= 2). Non-linear dimensional reduction was carried out by RunUMAP() function with dimensions 1:30. Each cluster was annotated by classical marker genes. Compositional differences in cell proportion between CBUs and non-CBUs were calculated by propeller method implemented in speckle v1.2.0 R package^33^. Differentially expressed genes between CBUs and non-CBUs were identified using the MAST test implemented in the FindMarkers() function, adjusting for cocaine use status. Only genes expressed in at least 10% of cells in either of the two groups were included in the analysis. Significance was defined by an FDR of less than 0.05 and an |avg_log2FC| exceeding 0.15. To explore the biological functions altered by long-term cannabis use, we performed Gene Ontology (GO) enrichment analysis for DEGs across cell types using R package clusterProfiler v4.10.1^64^. Benjamini-Hochberg was used to adjust P values, and the threshold was set to FDR<0.05. The top 30 enriched GO terms per cell type were displayed to facilitate interpretation of functional differences.

### Single nucleus ATACdata analysis workflow

Chromatin data was processed by Signac v1.14.0^38^ which interfaces seamlessly with the Seurat package^32^. After quality control, the filtered snATAC data was normalized with term frequency-inverse-document-frequency (TF-IDF) normalization. We then ran singular value decomposition (SVD) on the TD-IDF matrix, followed by batch effect correction with harmony v1.2.3^63^. Next, we run FindNeighbors() function with dimensions 2:50 and identified cell clusters by SLM algorithm in using FindClusters() at default resolution. The RunUMAP() function placed the similar cells together into a low-dimensional space with dimensions 2:50. Cell labels were transferred form RNA modality since snRNA and snATAC sequencing were profiled on the same cell population, the resulting datasets share identical cell barcodes. Then we generated UMAPs based on snRNA, snATAC, as well as combined data by integrating both RNA and ATAC data using WNN analysis^65^. Cell type-specific ATAC peaks were called using the MACS2 software (v2.2.9.1)^66^, which was invoked through the Signac package^38^ with default parameters for each major cell type. The FeatureMatrix() function was used to quantify peak count and created a MACS2 peak matrix. The cell type-specific peak matrix underwent reprocessing with TF-IDF normalization, SVD, and batch effect correction followed by dimension reduction as noted above. To evaluate differences in chromatin accessibility between CBUs and non-CBUs, the FindMarkers() function was applied to peaks present in at least 10% of cells in either group, using a logistic regression framework. The model was adjusted for total number of fragments and cocaine use status. Adjusted P values were used to determine significance at an FDR<0.05 and |avg_log2FC| cutoff was set as 0.15. ChIPseeker (v1.38.0) was used to annotate genomic regions and proximal genes corresponding to DARs.

### Identification of candidate *cis*-regulatory elements for DEGs

To investigate overall *cis*-regulatory architecture, the co-accessibility of genomic regions was assessed by R package cicero v1.3.9^67^. We computed correlations of chromatin accessibility of each genomic region pairs to identify putative enhancer-promoter interactions within each cell type. The furthest distance between two sites is 500kb and a 0.01 co-accessibility threshold was set. To gain deeper insights into the regulatory role of chromatin status in gene expression, we established connections by calculating the expression-accessibility correlation separated by major cell type. Firstly, we calculated Pearson correlation between expression of DEGs and accessibility of peaks overlapping their promoters (3kb upstream and downstream of TSS). Similarly, for potential distal regulatory elements of a given DEG, we selected peaks which are co-accessible with peaks coinciding with its promoter and positioned 3∼500kb from its TSS. Pearson correlation between gene expression and accessibility of the candidate enhancer was computed. cCRE links with absolute value correlation coefficient >0.05 and FDR<0.05 were retained. Only cCRE-peaks that were designated as DAR were kept.

### Differential transcription factor binding motif activity analysis

A list of motif position frequency matrices was retrieved from the JASPAR2020 database using the TFBSTools package (v1.40.0). To explore TFs that potentially play important roles associated with chronic cannabis use, an activity score was first estimated by RunchromVAR() function for each motif. Then differentially active motifs between CBUs and non-CBUs were subsequently detected by logistic regression model implemented in FindMarkers() function, accounting for sequencing depth and cocaine use status. Significant results were defined using a threshold of FDR<0.05.

### Transcription factor-target gene regulatory network

Through joint analysis of snATAC-seq and snRNA-seq data, we identified candidate target genes and construct cell-type-specific TF regulatory networks for selected TFs. For a given TF, candidate target genes were defined as genes whose promoters or distal cCREs contain binding motifs of this TF, and whose expression levels are significantly correlated with TF motif activity (FDR<0.05). Then we keep gene-TF links whose Pearson correlation coefficients fall within the top 5%. Cell type-specific TF regulatory networks were build using the R package igraph v2.1.3.

### Cell-cell communication analysis

CellChat v2.2.0 was used to infer cell-cell communication between major cell types based on the expression levels of ligands and receptors. By detecting overexpression of ligands or receptors in a cell group, CellChat infers ligand-receptor interactions when at least one component is overexpressed. Biologically significant intercellular interactions were inferred by assigning a probability value to each using the computeCommunProb() function, with parameters set to type=“truncatedMean” and trim=0.1. Differential cell-cell crosstalk between the two conditions were then identified and differential intercellular interaction network was constructed. First, we identified cell populations showing differences in overall interaction strength. Next, we determined the altered signaling which may drive communication. Ligand-receptor pairs with increased or decreased interaction probabilities were determined by subtracting those of the non-CBU group from the CBU group, and the expression levels of ligands and receptors were further examined.

### R package for visualization

BioRender (https://www.biorender.com/) were used to generate schematic diagram of the research framework. UMAPs, gene expression heatmaps, cell proportion bar plot were plotted by SCP v0.5.6. UpSetR v1.4.0 was used to generate the UpSet plots. scRNAtoolVis v0.1.0 was employed to draw volcano plot for DEGs of each cell type. Coverage and DNA sequence motif plots were generated by Signac v1.14.0., while TF regulatory networks were build using the R package igraph v2.1.3. Cell communication circle plot, heatmaps, pathway networks and interaction scatter plot were plotted by CellChat v2.2.0. Other figures were generated using ggplot2 v3.5.1, unless stated otherwise.

## Declarations

### Ethics approval and consent to participate

The study was approved by the committee of Human Research Subject Protection at Yale University and the Institutional Research Board Committee of the Connecticut Veteran Healthcare System. Informed consent was provided by all MWCCS participants via protocols approved by institutional review committees at each affiliated institution.

### Availability of data and materials

Data in the application part were from the MWCCS cohort, which has been identified as one with multiple vulnerabilities (e.g., racial/ethnic minority women, coinfected). Whereas participants from the cohort who contributed to the findings summarized in this manuscript provided written consent for genetic studies, said consent was collected prior to the most recent guidelines and requirements for data sharing. The MWCCS cohort operates under an alternative data sharing plan registered with the National Institutes of Health and access to data can be requested by submitting a Concept Sheet, which can be found along with instructions for Concept Sheet submission, at https://statepi.jhsph.edu/mwccs/.

The accession number for the WIHS in dbGaP genomic data is now provided (phs001503). The cohort is currently being re-approached to obtain informed consent for sharing of their data. This has been consistent with other genomic studies in the WIHS cohort. The algorithm for HBI and codes for analysis are on GitHub.

### Authors’ contributions

M. L. conducted data processing and analysis, contributed to interpretation of findings and manuscript preparation. K. A. performed data generation. X. D. contributed to data analysis. G. P. P. contributed to the interpretation of fundings. Y. H. participated in analytical strategies and supervision of data analysis. C. M. contributed to interpretation of the findings and manuscript preparation. M. H. C. collected the data. N. M. A. involved in data collection and interpretation, manuscript preparation. A. V. contributed to data analysis. D. H. and E. J. participated in study design, data interpretation, and manuscript preparation. B. E. A. oversaw the study, including study design, interpretation of the findings and manuscript preparation. K. X. oversaw the study, including study design, data processing, statistical analyses, interpretation of the findings, and manuscript preparation.

## Acknowledgements

The authors appreciate the support of the WIHS cohort and Yale Center of Genomic Analysis. Data in the application part of this manuscript were collected by the Women’s Interagency HIV Study (WIHS), now the MACS/WIHS Combined Cohort Study (MWCCS). The contents of this publication are solely the responsibility of the authors and do not represent the official views of the National Institutes of Health (NIH). MWCCS (Principal Investigators): Atlanta CRS (Ighovwerha Ofotokun, Anandi Sheth, and Gina Wingood), U01-HL146241; Baltimore CRS (Todd Brown and Joseph Margolick), U01-HL146201; Bronx CRS (Kathryn Anastos, David Hanna, and Anjali Sharma), U01-HL146204; Brooklyn CRS (Deborah Gustafson and Tracey Wilson), U01-HL146202; Data Analysis and Coordination Center (Gypsyamber D’Souza, Stephen Gange and Elizabeth Topper), U01-HL146193; Chicago-Cook County CRS (Mardge Cohen and Audrey French), U01-HL146245; Chicago-Northwestern CRS (Steven Wolinsky, Frank Palella, and Valentina Stosor), U01-HL146240; Northern California CRS (Bradley Aouizerat, Jennifer Price, and Phyllis Tien), U01-HL146242; Los Angeles CRS (Roger Detels and Matthew Mimiaga), U01-HL146333; Metropolitan Washington CRS (Seble Kassaye and Daniel Merenstein), U01-HL146205; Miami CRS (Maria Alcaide, Margaret Fischl, and Deborah Jones), U01-HL146203; Pittsburgh CRS (Jeremy Martinson and Charles Rinaldo), U01-HL146208; UAB-MS CRS (Mirjam-Colette Kempf, Jodie Dionne-Odom, Deborah Konkle-Parker, and James B. Brock), U01-HL146192; UNC CRS (Adaora Adimora and Michelle Floris-Moore), U01-HL146194. The MWCCS is funded primarily by the National Heart, Lung, and Blood Institute (NHLBI), with additional co-funding from the Eunice Kennedy Shriver National Institute Of Child Health & Human Development (NICHD), National Institute On Aging (NIA), National Institute Of Dental & Craniofacial Research (NIDCR), National Institute Of Allergy And Infectious Diseases (NIAID), National Institute Of Neurological Disorders And Stroke (NINDS), National Institute Of Mental Health (NIMH), National Institute On Drug Abuse (NIDA), National Institute Of Nursing Research (NINR), National Cancer Institute (NCI), National Institute on Alcohol Abuse and Alcoholism (NIAAA), National Institute on Deafness and Other Communication Disorders (NIDCD), National Institute of Diabetes and Digestive and Kidney Diseases (NIDDK), National Institute on Minority Health and Health Disparities (NIMHD), and in coordination and alignment with the research priorities of the National Institutes of Health, Office of AIDS Research (OAR). MWCCS data collection is also supported by UL1-TR000004 (UCSF CTSA), UL1-TR003098 (JHU ICTR), UL1-TR001881 (UCLA CTSI), P30-AI-050409 (Atlanta CFAR), P30-AI-073961 (Miami CFAR), P30-AI-050410 (UNC CFAR), P30-AI-027767 (UAB CFAR), P30-MH-116867 (Miami CHARM), UL1-TR001409 (DC CTSA), KL2-TR001432 (DC CTSA), and TL1-TR001431 (DC CTSA).

The authors gratefully acknowledge the contributions of the study participants and dedication of the staff at the MWCCS sites.

