## Supplementary figure and legend for "A single-nucleus multiome analysis of transcriptome and chromatin accessibility reveals cell-type-specific immune modulation for chronic cannabis use among people with HIV infection"

[2]VA Connecticut Healthcare System, West Haven, CT, U.S.A.

[3]Biomedical Informatics and Data Science, Yale School of Medicine, New Haven, CT, U.S.A.

[4]Translational Research Center, College of Dentistry, New York University, New York, NY, U.S.A.

[5]Department of Oral and Maxillofacial Surgery, College of Dentistry, New York University, New York, NY, U.S.A.

[6]Solutions Unit, RTI International, Research Triangle Park, NC, U.S.A.

[7]Fellow Program, RTI International, Research Triangle Park, NC, U.S.A.

[8]Center for Biomedical Information and Information Technology, National Cancer Institute, Rockville, MD, U.S.A.

[9]Miller School of Medicine, Division of Cardiovascular Medicine, University of Miami, Miami, FL, U.S.A.

[10]Department of Medicine, Stroger Hospital, Cook County Health System, Chicago, IL, U.S.A.

[11]UNC HIV Cure Center, University of North Carolina at Chapel Hill School of Medicine, Chapel Hill, NC, U.S.A.

[12]Department of Medicine, Division of Infectious Diseases, University of North Carolina at Chapel Hill School of Medicine, Chapel Hill, NC, U.S.A.

**Corresponding address to**

**Ke Xu, MD, PhD**

Professor of Psychiatry

Yale School of Medicine

Connecticut VA Healthcare

Biomedical Informatics and Data Science

300 George st, Suite 901

New Haven, CT 06511

### Supplementary Figures

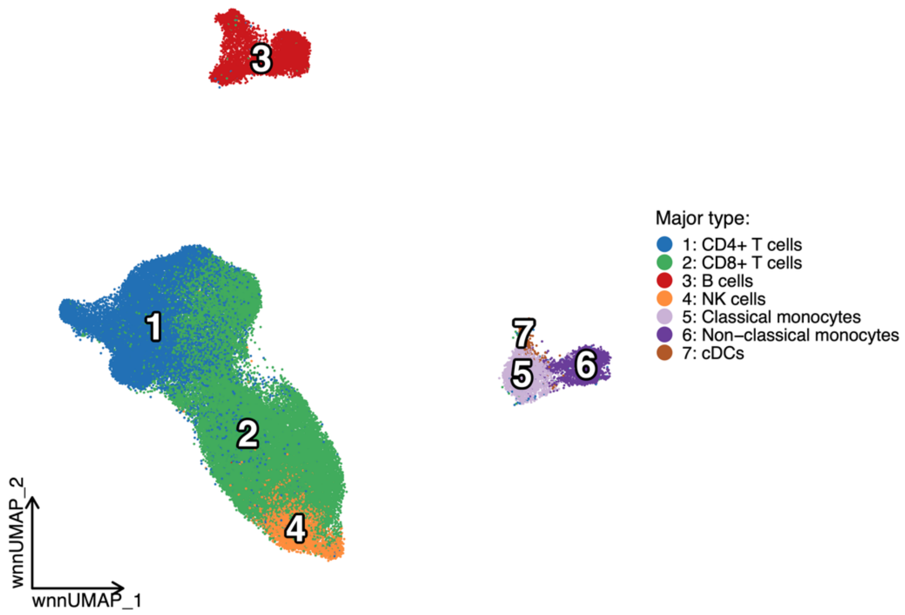


**Supplementary Figure 1. A UMAP showing major cell types of 80,480 nuclei from PBMC samples based on WNN-integrated data.** Each dot represents a nucleus and is colored according to major cell type. NK, natural killer cells; cDCs, conventional dendritic cells; WNN: weighted-nearest neighbor.

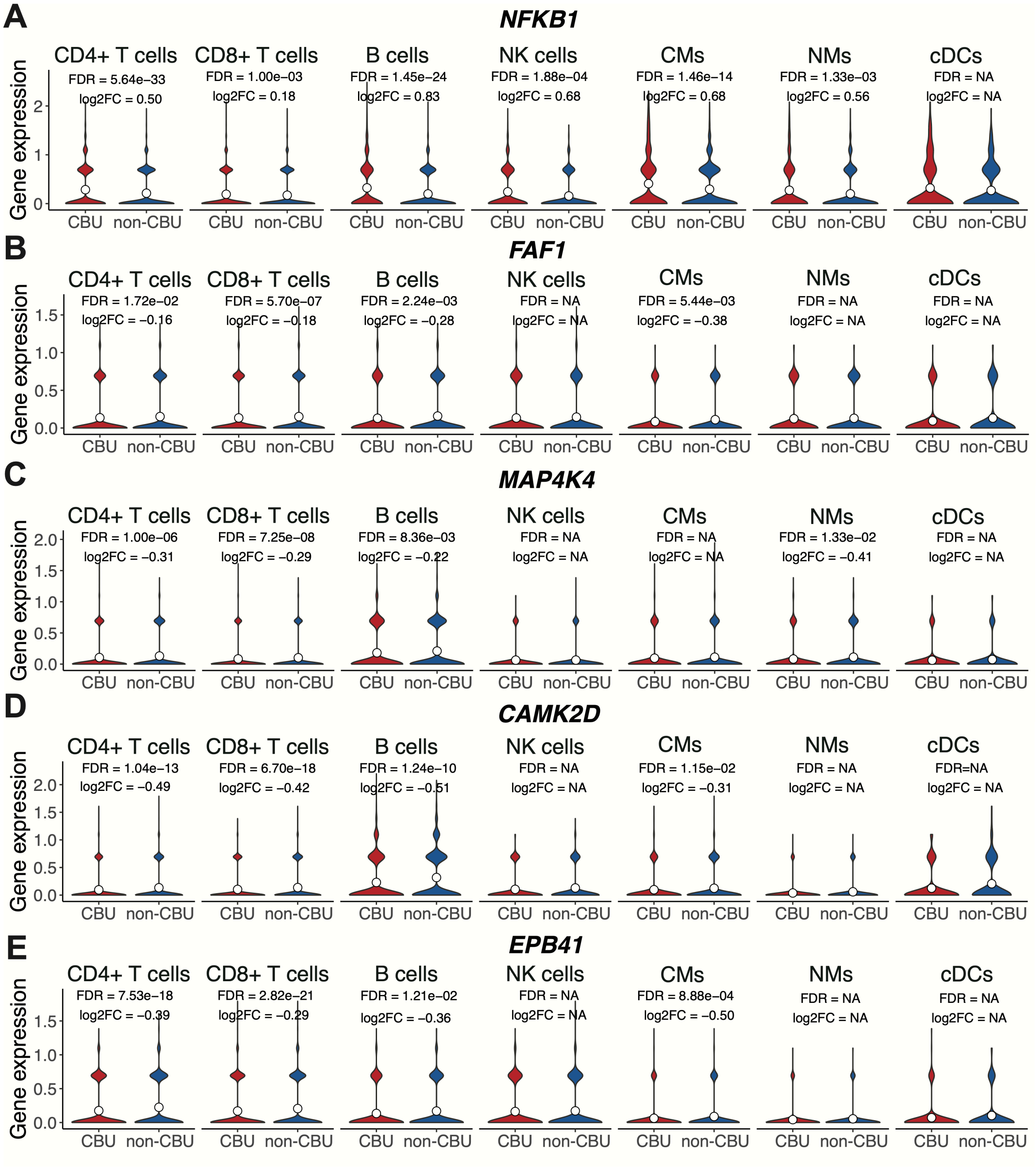


**Supplementary Figure 2. Shared Differentially Expressed Gene (DEG) expression across major cell types.** (A-E) Gene expression level of *NFKB1*; *FAF1*; *MAP4K4*; *CAMK2D*; *EPB41* across cell types in cannabis users (CBUs) and non-users (non-CBUs). For genes that are not differentially expressed in a given cell type, the FDR and log2FC are recorded as NA. FDR, false discovery rate; FC, fold change. NK cells, natural killer cells; CMs, classical monocytes; NMs: non-classical monocytes; cDCs, conventional dendritic cells.

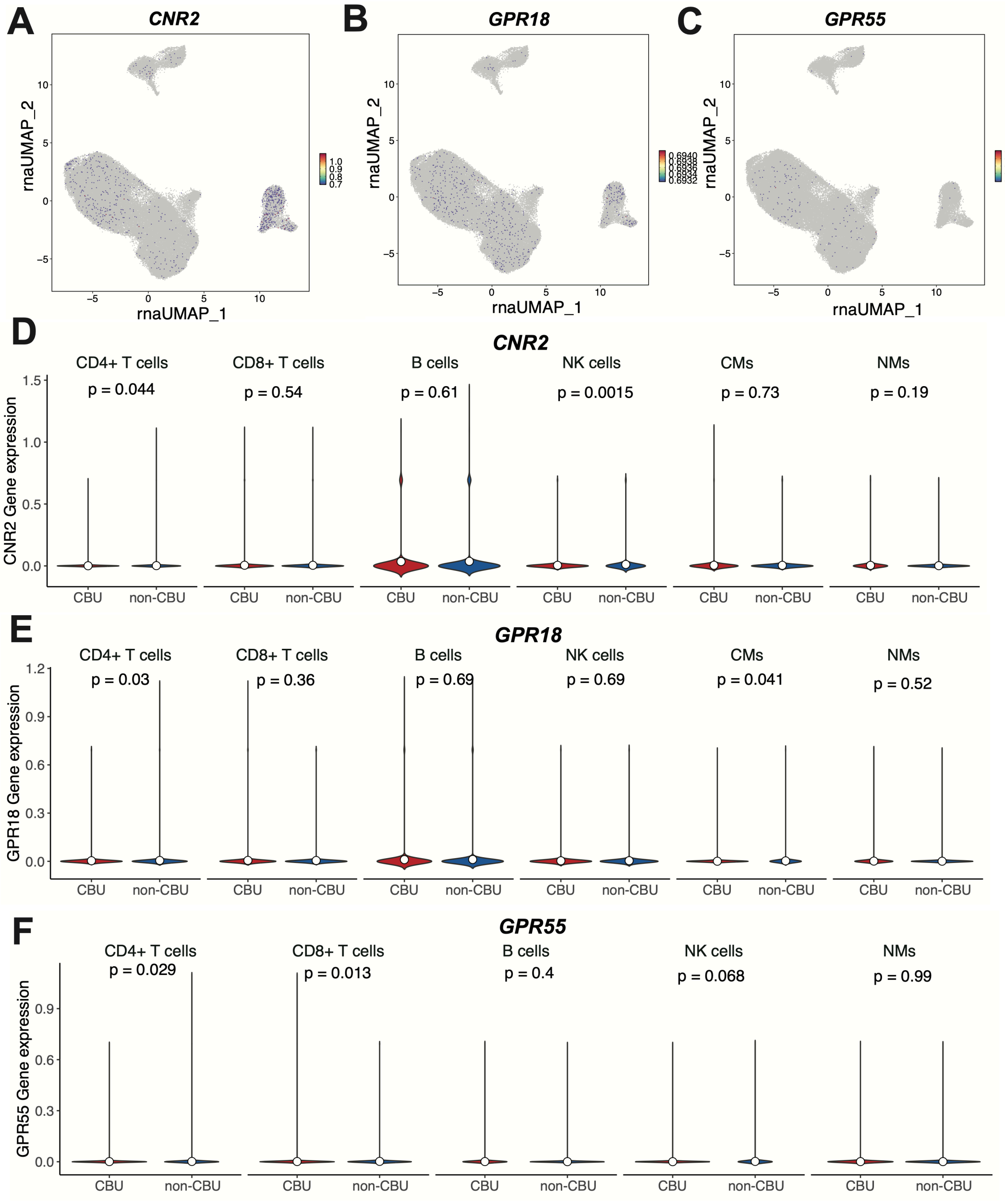


**Supplementary Figure 3. Expression of cannabis receptor across cell types.** (A-C) UMAPs and (D-F) violin plots showing the expression of *CNR2*, *GPR18*, *GPR55* across cell types. Violin plots in classical monocytes (CMs) and/or conventional dendritic cells (cDCs) were removed because genes are not expressed in these cell types. NK cells, natural killer cells; NMs: non-classical monocytes.

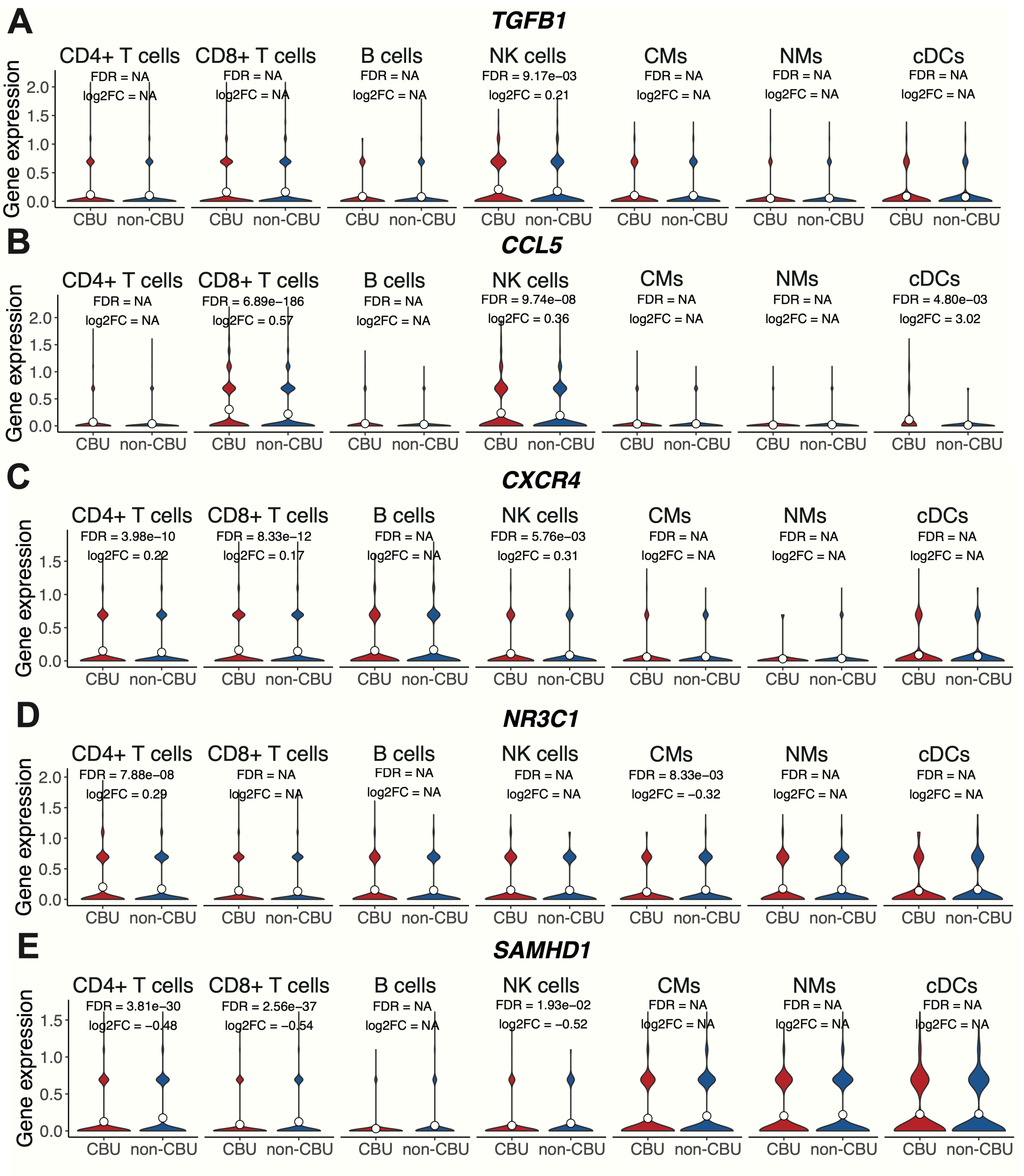


**Supplementary Figure 4. Expression of inflammation Differentially Expressed Gene (DEG) across all cell types.** (A-E) Gene expression level of *TGFB1*; *CCL5*; *CXCR4*; *NR3C1*; *SAMHD1* across cell types in cannabis users (CBUs) and non-users (non-CBUs). FDR, false discovery rate; FC, fold change. NK cells, natural killer cells; CMs, classical monocytes; NMs: non-classical monocytes; cDCs, conventional dendritic cells.

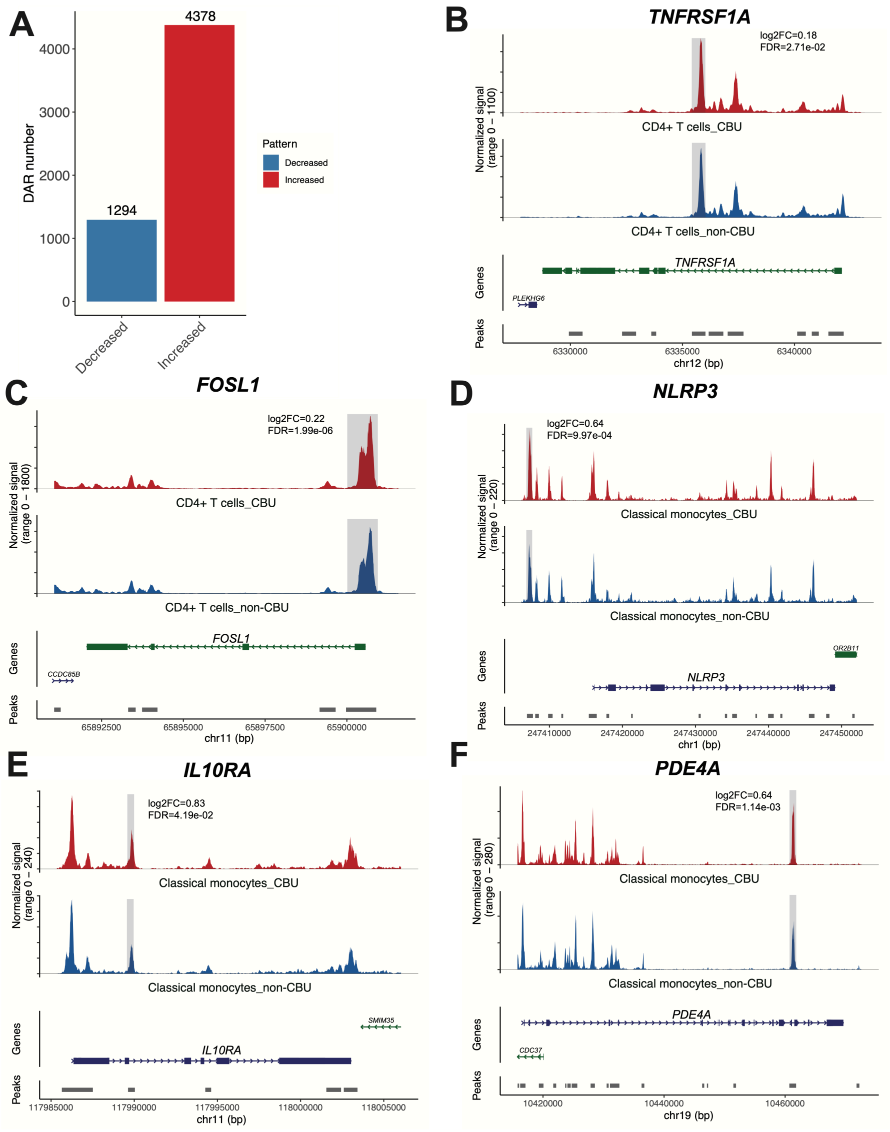


**Supplementary Figure 5. Differentially Accessible Regions (DARs) identified in CD4+ T cells and classical monocytes.** (A) A Bar pot demonstrating the number of DARs with increased or decreased accessibility. (B-F) Coverage plots illustrating examples of DAR and their proximal genes in specific cell type. (B and C) DAR near (B) *TNFRSF1A* in CD4+ T cells; (C) *FOSL1* in CD4+ T cells and (D-F) DAR proximal to (D) *NLRP3* in classical monocytes; (E) *IL10RA* in classical monocytes; (F) *PDE4A* in classical monocytes. The top panel shows chromatin accessibility, with cannabis user (CBU) shown in red and non-user (non-CBU) in blue. The bottom panel displays gene and peak annotations. DAR was highlighted with gray background. FDR, false discovery rate; FC, fold change.

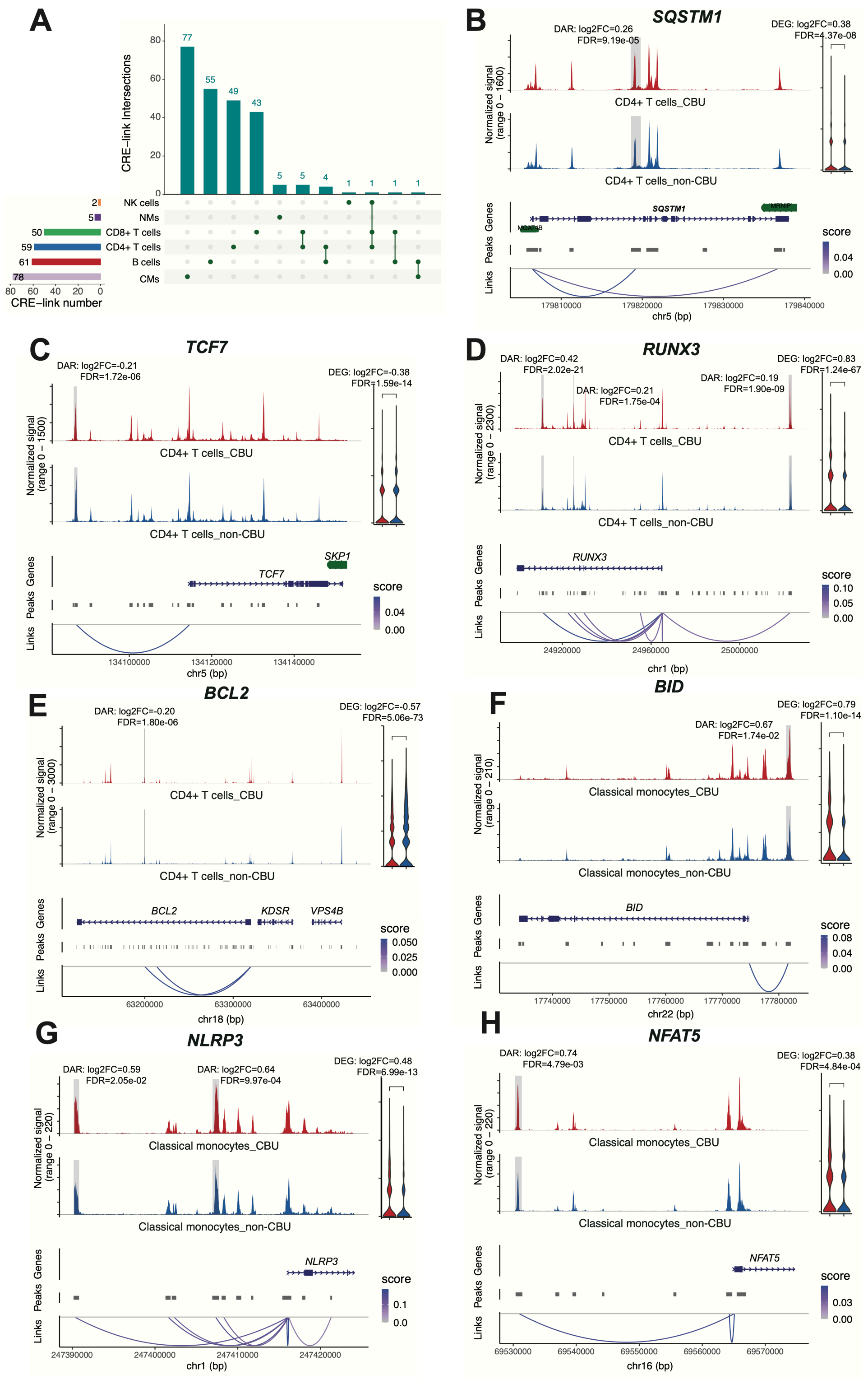


**Supplementary Figure 6. Cell-type-specific candidate *cis*-regulatory elements (cCREs) of Differentially Expressed Genes (DEGs).** (A) An upset plot showing the number of unique or shared cCRE-gene links across all cell types. (B-E) Coverage plots demonstrating cCRE-DEG links of (B) *SQSTM1* in CD4+ T cells; (C) *TCF7* in CD4+ T cells; (D) *RUNX3* in CD4+ T cells; (E)

*BCL2* in CD4+ T cells, and (F-H) cCRE-DEG links of (F) *BID* in classical monocytes; (G) *NLRP3* in classical monocytes*;* (H) *NFAT5* in classical monocytes. The top panel shows chromatin accessibility, with cannabis user (CBU) shown in red and non-user (non-CBU) in blue. The area of DARs significantly correlated with DEGs is highlighted in gray. The second panel displays the gene and peak coordinates. And the bottom panel show the peak-to-gene link between DEG and DAR. FDR, false discovery rate; FC, fold change.

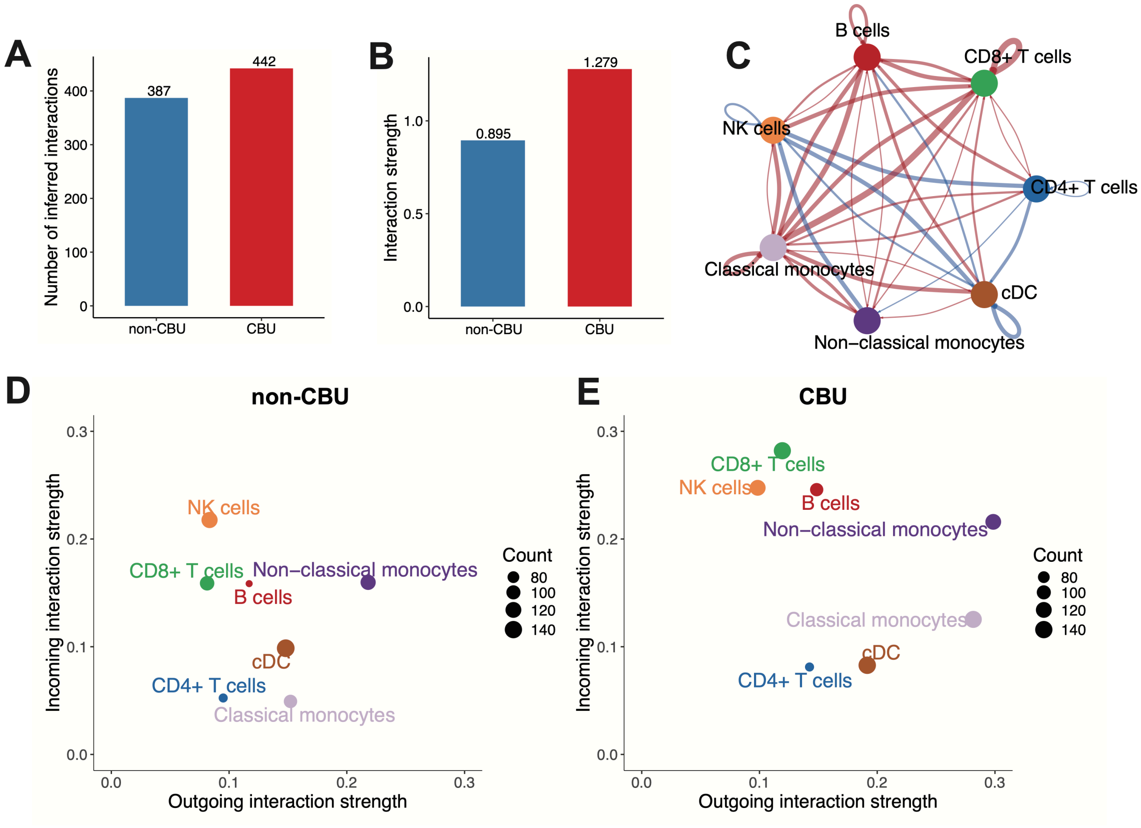


**Supplementary Figure 7. Cell-cell communication alteration among cell populations by cannabis use.** Bar plots showing the (A) total number of interactions; and (B) interaction strength in cannabis users (CBUs) and non-users (non-CBUs). (C) A circle plot showing differential number of interactions among major cell types. (D-E) Scatter plots showing cell populations with significant changes in sending or receiving signals in (D) non-CBU and (E) CBU group. NK cells, natural killer cells; CMs, classical monocytes; NMs: non-classical monocytes; cDCs, conventional dendritic cells.
